# Functional and structural properties of highly responsive somatosensory neurons in mouse barrel cortex

**DOI:** 10.1101/789347

**Authors:** C Barz, PM Garderes, D Ganea, S Reischauer, D Feldmeyer, F Haiss

## Abstract

Sparse population activity is a hallmark of supra-granular sensory neurons in neocortex. The mechanisms underlying sparseness are not well understood because a direct link between the neurons activated *in vivo* and their cellular properties investigated *in vitro* has been missing. We used two-photon calcium imaging to identify a subset of neurons in layer L2/3 (L2/3) of mouse primary somatosensory cortex that are highly active following principal whisker vibrotactile stimulation. These high responders were then tagged using photoconvertible green fluorescent protein for subsequent targeting in the brain slice using intracellular patch-clamp recordings and biocytin staining. This approach allowed us to investigate the structural and functional properties of high responders that distinguish them from less active control cells. Compared to less responsive L2/3 neurons, high responders displayed increased levels of stimulus-evoked and spontaneous activity, elevated noise and spontaneous pair-wise correlations, and stronger coupling to the population response. Intrinsic excitability was reduced in high responders, while other electrophysiological and morphological parameters were unchanged. Thus, the choice of which neurons participate in stimulus encoding may largely be determined by network connectivity rather than by cellular structure and function.

## Introduction

The perception of the world around us with its rich sensory information depends on the selective activation of neurons in the brain. While the activation of too many neurons is associated with cognitive impairment and pathological states like epilepsy, the activation of too few neurons can lead us to miss critical information from our environment. When neurons are activated, they fire action potentials, or spikes, which represent the primary mechanism of information transmission in the brain. How and why neurons are recruited into active spiking ensembles is still an open question, however. Two-photon imaging provides a key method to observe large populations of identified active and silent neurons, thereby enabling us to shed light on the principles of neuronal coding.

Mounting evidence indicates that sensory information is encoded by only a small number of active neurons in neocortex (Olshausen and Field 2004; Sachdev et al. 2012; Wolfe et al. 2010). This sparse coding scheme is a hallmark of L2/3 excitatory neurons in the primary somatosensory (barrel) cortex of rodents, where a small fraction of excitatory neurons has been found to fire the majority of single whisker stimulation evoked spikes (Crochet et al. 2011; de Kock et al. 2007; Kerr et al. 2007; Margolis et al. 2012; O’Connor et al. 2010).

Why only a subset of neurons is activated under these circumstances is a question under active investigation. It has been suggested that such sparseness may arise either from high trial-to-trial variability in the fraction of activated neurons or from a small subset of neurons that is consistently recruited by sensory stimulation (Barth and Poulet 2012). The latter possibility has been supported by the finding that a distinct set of L2/3 neurons is consistently activated by passive whisker stimulation over the course of days and months (Margolis et al. 2012; Mayrhofer et al. 2015b). In addition, sparseness might be related to specific stimulus parameters as other studies have described more variable responses (Kerr et al. 2007; Ranjbar-Slamloo and Arabzadeh 2017).

If specific stimuli activate only a small subset of neurons, it is possible that these cells show specific features in their morphology (Oberlaender et al. 2012), physiological cell properties (Crochet et al. 2011) or connectivity (Benedetti et al. 2013) that make them more responsive compared to other neurons. However, studies that have attempted to address these questions have failed to provide a direct link between the neurons activated *in vivo* by whisker stimulation and the cellular and circuit properties investigated *in vitro* (Elstrott et al. 2014; Yassin et al. 2010). To overcome this limitation, we have chosen a combination of *in vivo* and *in vitro* methods (Fig. 1): We used two-photon calcium imaging to identify neurons in L2/3 of the mouse primary somatosensory cortex that are strongly activated by vibrotactile stimulation of the principal whisker. The individual high responders were then tagged using light activation of the photo-convertible green fluorescent protein (PAGFP; Lien and Scanziani 2011). The tagged neurons showed strong fluorescence so that they could be identified in the brain slice and selected for intracellular patch-clamp recordings and biocytin stainings (Marx et al. 2012). The combination of techniques made it possible to perform a detailed characterization of identified high responders and their physiological and morphological properties that may explain their elevated activity levels. During *in vivo* imaging, high responders (HRs) exhibited strong, fast and reliable activation by whisker stimulation as well as increased levels of spontaneous activity. Compared to less active neurons, HR activity was more correlated during stimulation and non-stimulation epochs and showed stronger coupling to the population response. Furthermore, *in vitro* intracellular measurements of tagged neurons revealed reduced intrinsic excitability as well as larger and more frequent excitatory network inputs for high responders compared to control neurons. Both groups displayed similar passive cell properties and similar morphological characteristics pertaining to different excitatory subclasses. Our findings support the view that a subset of excitatory L2/3 neurons are preferentially targeted by excitatory network inputs, leading to their increased activation both in the presence and absence of sensory stimulation and to a compensatory down-regulation of intrinsic excitability in this population. We conclude that the choice of which neurons participating in stimulus coding may largely depend on local network connectivity, rather than on the specific morphological and physiological characteristics of the neurons.

**Fig. 1.**
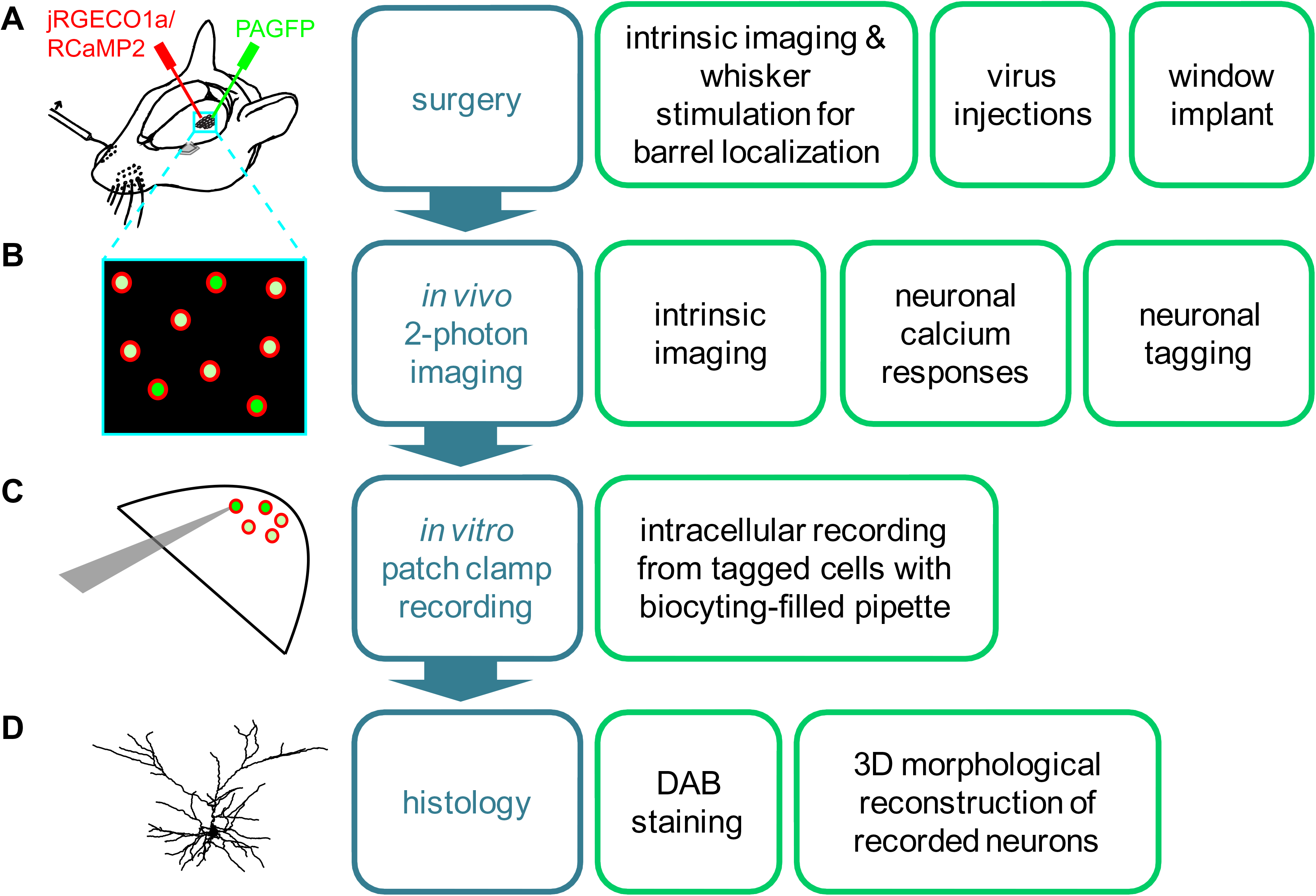
Combined *in vivo* and *in vitro* approach for the functional and structural characterization of neurons with known stimulus-response properties. The method comprised four main experimental steps (*A-D*): **(A)** Intrinsic imaging for localizing whisker-evoked response and injection site, followed by injections of two viruses and a chronic window implant. The viruses comprised a red calcium indicator (jRGECO1a or RCaMP2) and a photo-activatable green fluorescent protein (PAGFP). **(B)** Two-photon imaging of whisker-evoked responses and neuronal tagging by light-activation of PAGFP-postive neurons. **(C)** Intracellular recordings of neurons tagged *in vivo* and simultaneous dye injection (biocytin). **(D)** Neuronal staining and reconstruction of neuronal morphology.

## Results

### High and low responding neurons identified by in vivo whisker stimulation and two-photon imaging

We characterized whisker-evoked responses in L2/3 neurons in mouse barrel cortex using two-photon imaging of a genetically encoded calcium indicator (jRGECO1a or RCaMP2; Fig. 2). L2/3 neurons were chosen based on a) the depth of their soma relative to the cortical surface (100-300 μm below the pia) and b) on the visual assessment of the density of cell bodies in L2/3, which is substantially higher compared to L1. In addition, targeted neurons in L2/3 could be visually distinguished from L4 neurons since neither of the two calcium indicators was expressed in L4.

**Fig. 2.**
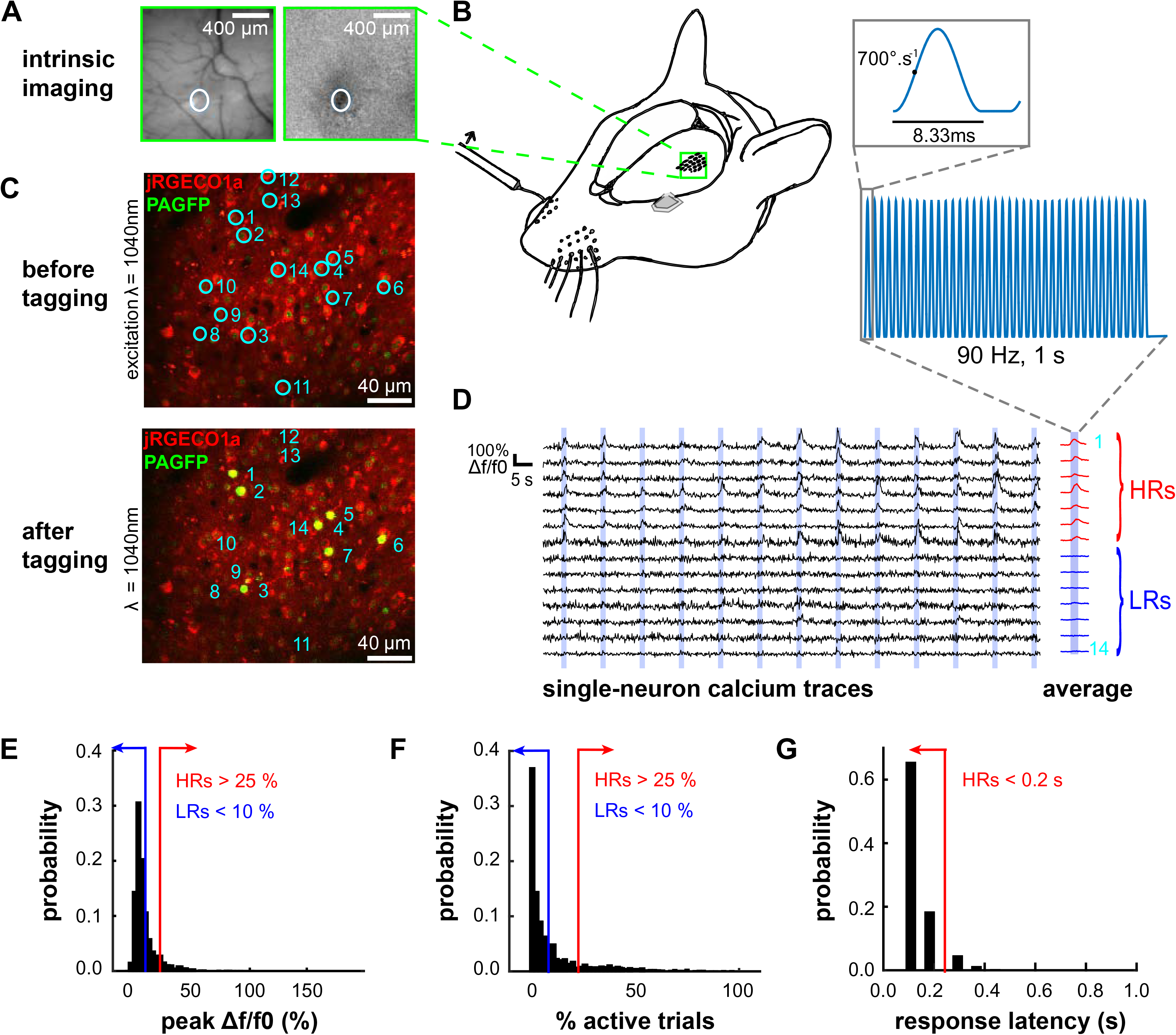
*In vivo* whisker stimulation, two-photon imaging and neuronal tagging. **(A)** Intrinsic imaging was performed prior to each two-photon imaging session to identify barrel location (white circle). Vasculature under green light illumination (*left*) and change in reflected red light RD/R0 during single whisker stimulation (*right*). **(B)** Mice (n = 30) were kept under light isoflurane anesthesia during two-photon imaging in barrel cortex L2/3 (green square; *left*) and piezo-electric stimulation of single whiskers at 90 Hz (*right*). **(C)** Co-expression of red calcium indicator for two-photon imaging (jRGECO1a or RCaMP2) and photo-activatable green fluorescent protein (PAGFP) for photo-tagging cells. Overlay of green and red channels before (*top*) and after photo-activation of PAGFP (*bottom*). **(D)** Single-neuron calcium transients Δf/f0 (black) in response to whisker stimulation (blue). High responders (HRs) #1-7 showed large average responses (red curves), while low responders (LRs) #8-14 showed small average responses (blue curves). **(E-G)** Criteria for defining HRs (red) and LRs (blue): Peak calcium signals (Δf/f0) averaged across all trials (*E*), fraction of trials with significant activation (*F*), and response latency (*G*). Neurons that did not match these criteria were categorized as medium responders.

To ensure that the imaging field of view (FOV) was centered on the target barrel, intrinsic optical imaging was performed prior to each two-photon imaging session (Fig. 2A). Depending on the area in which the calcium indicator was expressed, we imaged between one and five barrels per animal. During imaging, the animal was kept under light isoflurane anesthesia and the principal whisker was stimulated with 1 s pulses at 90 Hz (Fig. 2B). This stimulation frequency was chosen since previous research has shown that a highly responsive subset of neurons can be reliably activated in the 10-110 Hz range and that spiking-related calcium signals are stronger at higher frequencies (Mayrhofer et al. 2015b).

Neurons were found to show various degrees of responsiveness to whisker stimulation – with some showing high levels of activity (Fig. 2C, top, and 2D, cells #1-7) and others low levels (cells #8-14). To define high and low responding neurons quantitatively, we used three criteria (Fig. 2E-G): First, neurons were defined as high responders (HRs) if they showed peak calcium signals (Δf/f0) at or above 25 % averaged across all trials. Second, HRs displayed significant activation relative to baseline in at least 25 % of the recorded trials. Third, response latencies of HRs did not exceed 200 ms following stimulus onset. In contrast, neurons were categorized as low responders (LRs) if their responses showed low amplitudes (≤ 10 % peak Δf/f_0_) and low reliability (≤ 10 % trials with significant activity), irrespective of latency.

Co-expression of a histone-bound, photo-activatable green fluorescent protein (PAGFP; Lien and Scanziani 2011) enabled us to selectively tag single neurons (Fig. 2C, bottom) for later identification in the brain slice and targeted intracellular recordings. Per animal, tagged neurons were classified as either HR or LR, but never mixed, so that either class could be identified unequivocally in the brain slice used in patch experiments.

### A gradient of neuronal activity in response to whisker stimulation

Supporting the notion of sparse coding in barrel cortex, only a small subset of highly active neurons was recruited during whisker stimulation, while the majority of neurons remained almost silent (Fig. 3A). The analysis included 2100 manually selected neurons that co-expressed the red calcium indicator (jRGECO1a or RCaMP2) and the PAGFP. Approximately 30 % of all neurons recorded showed no activity in any trial. By applying our criteria, we categorized neurons as HRs (Fig. 3A, top row, red lines) and LRs (blue lines). Neurons that did not fit these categories were classified as medium responders (MRs; grey lines), displaying an intermediate level of responsiveness. On average, HRs and LRs showed large and small response amplitudes, respectively (Fig. 3B). Taking a closer look at the distribution of response amplitudes, latency and fraction of trials active, a gradient of responses emerged with HRs and LRs at the opposite ends of the spectrum (Fig. 3C). This finding supports earlier results that neurons with high and low levels of activity are not separate cell clusters, but rather two extremes observed in the neuronal population (Elstrott et al. 2014).

**Fig. 3.**
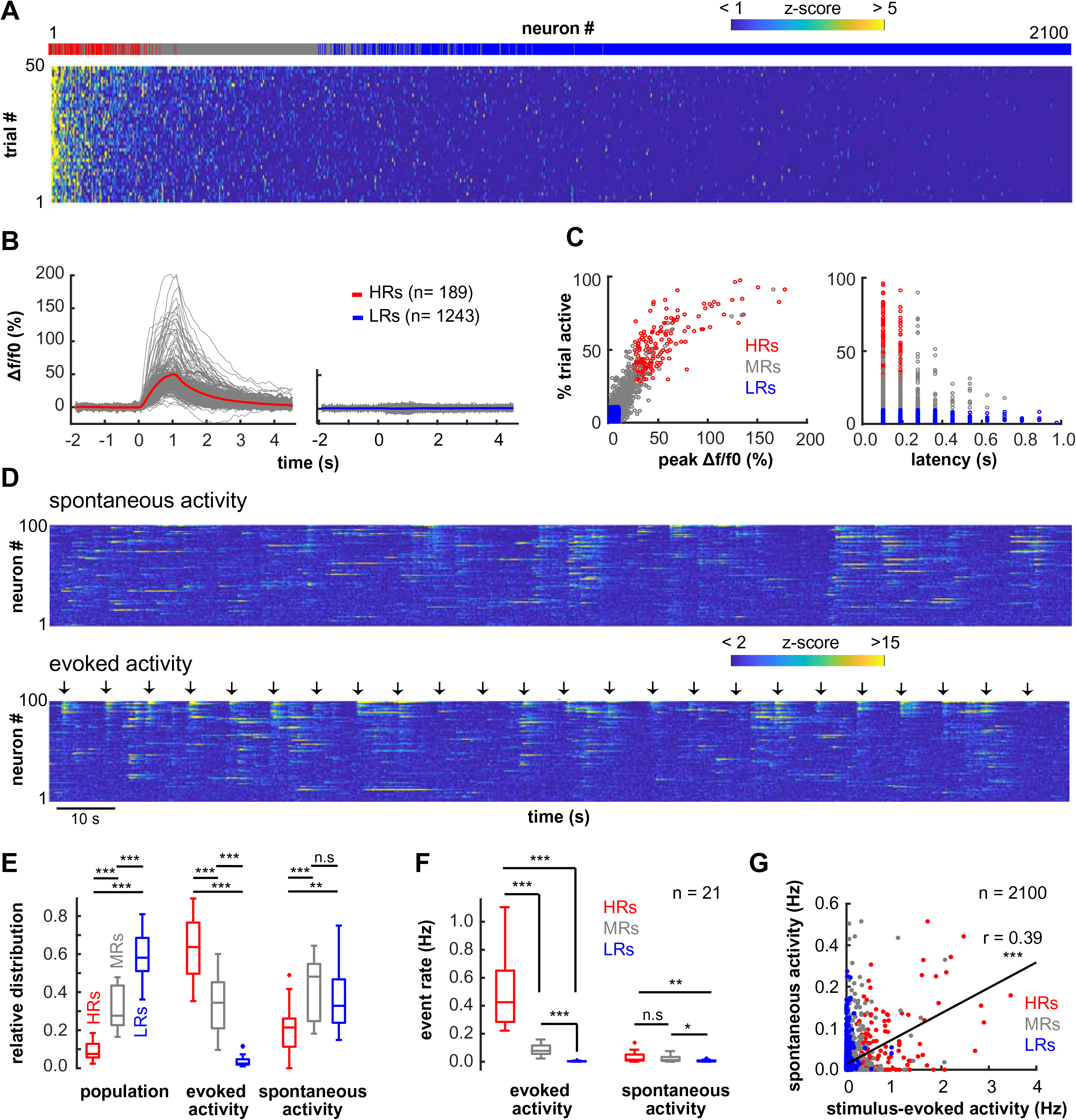
A gradient of neuronal activity in response to whisker stimulation. High responders (HRs; n = 189), low responders (LRs; n = 1243) and medium responders (MRs; n = 668) were recorded in 21 field of views (FOVs; n = 21 mice). **(A)** Neuronal calcium responses during trials, ranked according to their average z-scores. Top row identifies neurons as HR (red), MR (grey) or LR (blue). **(B)** Response amplitudes of HRs (*left*) and LRs (*right*). **(C)** Trial-to-trial variability as a function of the stimulus-evoked peak amplitude *(left)* and the response latency (*right*). Each dot represents one neuron. **(D)** Discrete events were extracted from continuous two-photon recordings using a deconvolution algorithm. Shown are z-scored events during spontaneous (*top*) and stimulus-evoked activity (*bottom*) for a representative set of 100 neurons recorded in one FOV. Each arrow indicates stimulus onset. **(E)** *Left-to-right:* Average distribution of HRs (red), MRs (grey) and LRs (blue) in the population (n = 2100 neurons, n = 21 FOVs), probability of calcium events during stimulation (n = 1504 trials) and during spontaneous activity (262 s on average). Statistical comparisons were performed using Mann-Whitney *U* tests. **(F)** Average event rates across cell classes during stimulus-evoked and spontaneous activity (Mann-Whitney *U* tests). **(G)** Relationship between the frequency of stimulus-induced and spontaneous calcium events across all neurons (Spearman rank correlation). Trend line (black) based on least squares fit. P-values: * < 0.05; ** < 0.01; *** < 0.001.

To assess the contribution of each responder category to the total population activity, we applied a deconvolution algorithm to the recorded calcium signals (see **Methods**) and extracted discrete spiking-related calcium events during stimulation that were significantly different from baseline levels (Fig. 3D). On average, stimulation increased the event rate in the entire neuronal population by a factor of three compared to baseline levels. Although HRs only represented a minority of neurons in the population, they contributed the majority of whisker-evoked spikes (Fig. 3E). In contrast, LRs constituted the majority of neurons in the population, but fired a minority of spikes in response to whisker stimulation. Again, MRs displayed an intermediate pattern of activity. Of note, these proportions were stable across FOVs and did not depend on the calcium indicator used (RCaMP2: mean ± SEM, 4.07 ± 0.88, n = 14; jRGECO1a: 4.69 ± 0.32, n = 71; p = 0.94; Mann-Withney *U* test). During epochs with spontaneous firing, HRs also contributed more spikes than one would expect based on their low prevalence in the population, while LRs contributed less than expected. With respect to average firing rates, HRs showed higher spontaneous and stimulus-evoked spiking activity than LRs (Fig. 3F). While average firing rates were higher during stimulation compared to spontaneous activity for HRs (Friedman test; p = 3.12e-08) and MRs (p = 3.68e-06), firing in LRs was similar in both conditions (p = 0.74). Furthermore, we found a positive correlation between stimulus-evoked and spontaneous spiking activity, with HRs scoring high on both measures, and MRs and LRs showing lower levels of activity in both conditions (Fig. 3G). These observations support earlier findings (Clancy et al. 2015; Kerr et al. 2007) and suggest that L2/3 neurons do not only show sparse coding, but more generally sparse population activity that is independent of stimulus parameters.

### High responders show elevated coupling to the network and to each other

While stimulation led to an overall increase in calcium responses in the neuronal population, we observed a large trial-to-trial variability in the ensemble size of the recruited neurons, *i.e*. the proportion of cells responding to stimulation in single trials. This proportion of recruited neurons ranged from 0 to 37.5 % of the total population. To explore the possibility that HR and LRs were recruited during different states of network activation, we analyzed their activity profiles in more detail (Fig. 4A-C). First, we tested whether random, independent firing of neurons in the FOV could account for the variability in responding ensemble size and shuffled the activity of each neuron across trials. (Kerr et al. 2007; see **Methods**). If single neurons were independent from overall network activity, shuffling the trials where they become active should not affect the trial-to-trial variability of the population response. Compared to the shuffled data, the actual experimental data was found to show a higher probability of trials with either very small responding ensembles (0-5 % of the total population) or very large responding ensembles (> 25 % of the total population; Fig. 4A). The discrepancy between shuffled and experimental data indicates that the recruitment of neurons did indeed covary with the network activity.

**Fig. 4.**
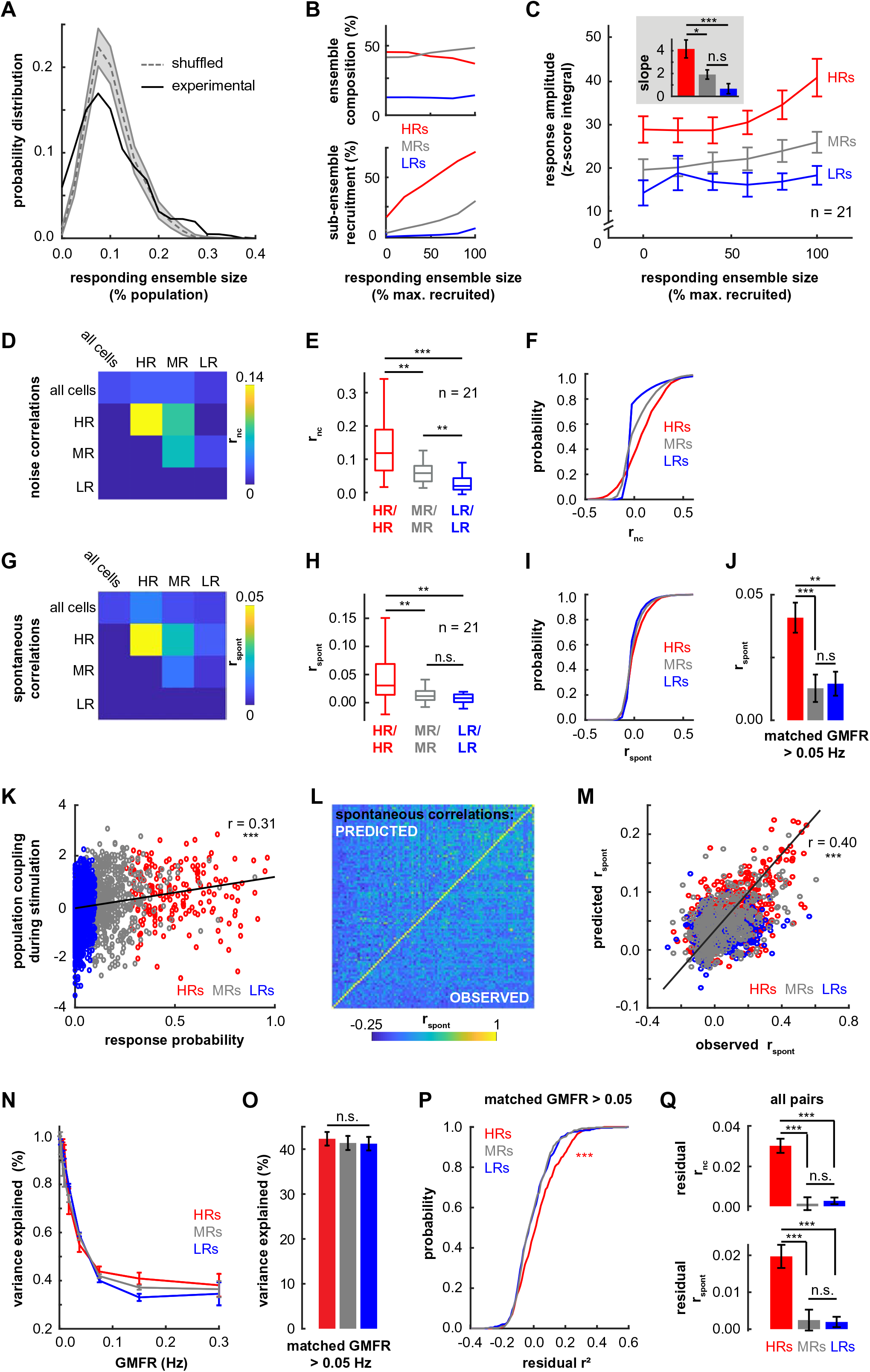
Network-dependent response profiles during stimulation and spontaneous activity. Data comprised n = 189 HRs, n = 668 MRs and n = 1243 LRs measured in 21 FOVs (n = 21 mice). Neuronal activity was analyzed with respect to changes in the fraction of active neurons (“ensemble size”, *A-C*), correlations among cell classes (*D-J*), and the degree of coupling to the population activity (*K-Q*). **(A)** Probability of ensemble activation (% of neural population activated by stimulus) under shuffled (dashed line) and experimental conditions (continuous line). For each neuron, activity was shuffled across trials while keeping the total activity constant. Data (n = 1504 trials) were pooled across FOVs (n = 21). Error shades represent shuffled data from 1^st^ to 99^th^ percentile. **(B)** Stimulus-induced changes in ensemble composition (*top*) and sub-ensemble recruitment (*bottom*) as a function of responding ensemble size. **(C)** Changes in response amplitude of different cell classes as a function of the size of the responding ensemble. The response amplitude was defined as the area under the curve of the z-scored calcium transient. The ensemble size was normalized by the maximum ensemble size per FOV across trials. *Inset:* Slope of the linear regression of the amplitude on the responding ensemble size. **(D-F)** Pair-wise correlations during stimulation. Correlations were quantified using Pearson r², which was averaged across 68 ± 10 trials for each FOV, and then averaged across FOVs for each cell class. Shown are the average r² between pairs of neurons during stimulation across FOVs (*D*), the corresponding boxplots with the median (midline), interquartile ranges and 2.7 SD (*E*), and the cumulative probability distributions of single pair-wise correlations with n = 1015 HRs/HRs, n = 11835 MRs/MRs, n = 38875 LRs/LRs (*F*). Statistical comparisons were carried out using Mann-Whitney *U* tests. **(G-J)** Pair-wise correlations during spontaneous activity. Pearson r² was calculated based on 258 ± 199 s of spontaneous activity. Displayed are the average r² between pairs of neurons during baseline across FOVs (*G*), the corresponding boxplots (*H*), and the cumulative probability distributions of single pair-wise correlations (*I*). Average spontaneous correlations for neuronal pairs (n = 487 HRs/HRs, n = 487 MRs/MRs, n = 487 LRs/LRs) with matched geometric mean firing rates (GMFR > 0.05 Hz) are shown in *J*. P-values: * < 0.05; ** < 0.01; *** < 0.001. **(K)** Population coupling as a function of response probability during stimulation. Across all neurons (n = 2100), population coupling and response probability were positively correlated (Spearman rank correlation). **(L)** Spontaneous correlations could be predicted by a simple model based on the mean event rate per neuron, the population rate per time bin, and the population coupling of the neuron. Shown are the pair-wise correlations for all pairs of neurons in an example FOV, with a comparison of the predicted (upper left triangle) and the experimentally observed pair-wise correlations (lower right triangle). **(M)** Pair-wise correlations predicted by the model as a function of experimentally observed correlations in 487 pairs of neurons with matching GMFR during spontaneous activity. **(N)** Prediction strength of the model across the three cell classes as a function of GMFR (spontaneous activity). **(O)** Variance in spontaneous correlations explained by the model for neurons with matching GMFR above 0.05 Hz (Mann-Whitney *U* test). **(P)** Cumulative distribution of residual correlations during spontaneous activity in HRs, MRs and LRs with matching GMFR above 0.05 Hz. **(Q)** Average residual correlations during stimulation (*top*) and spontaneous activity (*bottom*) for all neuronal pairs (Mann-Whitney *U* test). Error bars represent 95 % confidence intervals (C, J, O, Q).

We then asked how the composition of the responding ensemble changed (*i.e*. which responder category was recruited) as a function of the size of the responding ensemble. Data from all FOVs were pooled after normalization to the most active ensemble recruited per FOV. Largely independent of the responding ensemble size, HRs and MRs dominated the population of activated neurons, while LRs contributed very little to the stimulus-evoked activity (Fig. 4B, top). In addition, we measured the fraction of HRs, LRs and MRs that were recruited as the size of the responding ensemble changed (Fig. 4B, bottom). We found a monotonic increase for all responder types, with HRs being recruited faster than the others. When the responding ensemble size was large, roughly 70 % of all HRs were active, while less than 10 % of the cells in the LR ensemble were active.

Knowing that the size of the responding ensemble is related to the types of responders that become activated, we asked whether ensemble size also affects the neurons’ response amplitudes. Indeed, we found that HRs increased their response amplitudes with increasing size of the responding ensemble (Fig. 4C). Compared to HRs, LR and MR cells showed significantly lower response amplitudes and lower dependence of the latter on network activation states (Fig. 4C, inset).

If network activity affects the recruitment of LRs and HRs, the question arises which neurons in the population tend to be activated together. To evaluate this, we computed pair-wise noise correlations and spontaneous correlations using Pearson‘s R². During stimulation, the highest average pair-wise correlations were measured among HRs, while correlations among LRs and MRs were less strong (Wilcoxon rank-sum test, p < 0.01; Fig. 4D-F). The analysis of average pair-wise correlations during epochs of spontaneous activity yielded the same conclusion (Fig. 4G-I). Since elevated spiking activity in HRs could partly explain higher levels of correlations in this group, we carried out a control analysis that only included pairs with matching geometric mean firing rates (GMFR; *i.e*. the average firing rate in any pair of neurons; see **Methods** & Cohen et al 2011). The results of this analysis confirmed increased levels of correlations among HRs relative to MRs and LRs (Fig. 4J). The finding of enhanced correlations in HRs during both stimulus-evoked and spontaneous epochs further supports the notion of stimulus-independent properties that differentiate high from low responding neurons.

It is conceivable that the increased correlations in HR activity result from shared, non-specific synaptic inputs from the local network. In line with this notion, a previous study in visual cortex showed that neurons with enhanced coupling to the firing of the population receive more excitatory inputs from neighboring neurons, independent of sensory tuning (Okun et al. 2015). Thus, population coupling was found to predict a large portion of spontaneous pair-wise correlations, and was a better predictor than stimulus tuning (Okun et al. 2015). Following the same approach, we first measured population coupling and found a positive correlation between population coupling and response probability (linear fit, r² =0.31, p <10^-16^). HRs scored highest on population coupling and response probability, while LRs were found at the opposite end of the spectrum (Fig. 4K). We then asked whether enhanced pair-wise correlations among HRs could result from stronger population coupling. We therefore compared the experimental data to a simple random generative model (Okun et al. 2015) that matched the experimental data in terms of the coupling of single neurons to the population, the mean firing rate of the neurons and the population rate (see **Methods**). During spontaneous network activity, the observed pair-wise correlations were predicted to a large degree by the model (Fig. 4L,M). More specifically, the model explained the largest variance in spontaneous correlations for neurons with low activity (GMFRs < 0.05 Hz; Fig. 4N). To avoid potential bias due to the dependence of the model’s performance on the firing rate, we repeated the analysis using pairs of neurons with matching GMFR. For neurons with a GMFR above 0.05 Hz, the model predicted approximately 40 % of the spontaneous correlations among HRs, MRs and LRs (Fig. 4O). After subtraction of predicted from observed spontaneous correlations, residuals for HRs remained significantly positive, while residuals for MRs and LRs were on average close to zero (Fig. 4P). This analysis was repeated for pairs of neurons during stimulus-evoked activity and the results showed again increased residuals for HRs (Fig. 4Q). This suggests that population coupling cannot fully explain the strength of functional connections in the population of HRs. Rather than being indistinct recipients of inputs from neighboring neurons, HRs appear to share more specific, one-to-one functional connections, thus creating a functional subnetwork.

### High responders show reduced intrinsic excitability

We first aimed to test whether L2/3 pyramidal neurons that were highly responsive to whisker stimulation had electrophysiological properties different from those of less responsive cells. Thus, we performed *in vitro* patch clamp recordings on cells previously imaged *in vivo*. Thalamocortical brain slices showed clear fluorescent PAGFP-labeling of neurons in layers 1, 2/3, 5 and 6 of the barrel cortex. PAGFP is bound to the histone protein H2B and is thus expressed in the nucleus (Lien and Scanziani 2011). As a result, tagged HRs and LRs appeared as distinctly bright spots in L2/3 and could be targeted for recordings (Fig. 5A,B).

**Fig. 5.**
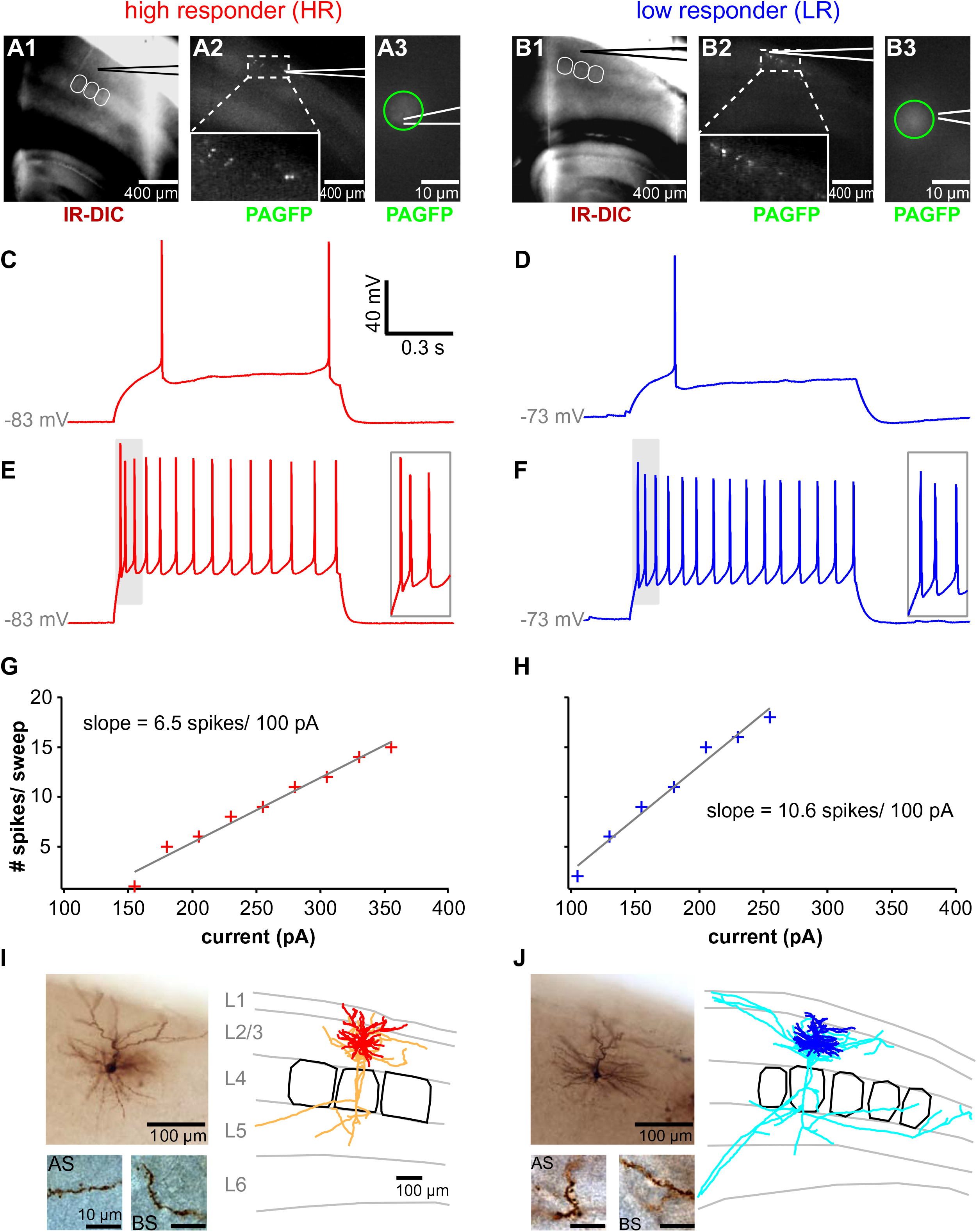
Electrophysiological and morphological characterization of high responders (HRs) and low responders (LRs) in barrel cortex L2/3. **(A)** Example of patched HR. Thalamocortical slice and pipette position under infrared-differential interference contrast (IR-DIC; *A1*) to visualize barrels (white), and under fluorescent illumination to visualize PAGFP-labeled cells (*A2*). HRs previously tagged *in vivo* appear brighter than non-tagged neurons (*A2, inset*). Fluorescent micrograph (*A3*) of patched neuron with PAGFP-labeled nucleus (green circle) and the biocytin-filled patch pipette (white line). **(B)** Example of patched LR. Thalamocortical slice (IR-DIC, *B1*) with brightly tagged LRs (PAGFP; *B2, B3*). Same conventions as in *A*. **(C,D)** Minimum current required to elicit spiking in HR (160 pA; *C*) and LR (100 pA; *D*). **(E,F)** Sustained firing pattern in HR (330 pA; *E*) and LR (205 pA; *F*). Insets display the first three spikes from the trace (grey shading). **(G,H)** Number of spikes fired as a function of current injected in HR (*G*) and LR (*H*). Data points are indicated by crosses and the fitting curve by a grey line. **(I,J)** Histological processing and morphological reconstruction of biocytin-filled HR (*I*) and LR (*J*). *Left panel:* Micrograph of neuron (*top*) with enlarged view of apical spines (AS; *bottom left*) and basal spines (BS; *bottom right*; scale bar: 10 µm). *Right panel:* 3D neuronal reconstruction with dendrites (HR: red, LR: blue), axons (HR: orange, LR: cyan), layer borders in grey and barrels in black. For 3D animations of the two example cells, see *Supplementary Materials*.

Based on previous findings (Elstrott et al. 2014; Yassin et al. 2010), we hypothesized that excitability is altered in the population of L2/3 excitatory HRs compared to LRs. Using *in vitro* patch clamp recordings and post-hoc histological reconstructions (see **Methods**), we identified excitatory neurons based on their firing pattern and morphological features. Among all tagged HRs, 76.2 % (16/21) were pyramidal neurons, while 23.8 % (5/21) were fast-spiking interneurons. Of the tagged LRs, 70.6 % (12/17) were pyramidal neurons, 17.6 % (3/17) were fast-spiking interneurons and 11.8 % (2/17) were non-fast spiking interneurons. Fig. 5 shows an example HR (left column) and LR (right column) with clear fluorescent labeling (**A,B**), typical firing properties (**C-H**) and classical pyramidal neuron morphology (**I,J**). To detect changes in excitability, we measured input resistance (R_input_), resting membrane potential (V_rest_), rheobase, firing frequency across steps of current injections, and spike threshold in 16 HR pyramidal neurons (n = 16 mice) and 12 LR pyramidal neurons (n = 9 mice). In addition, we quantified the initial doublet spike burst as the first inter-spike interval (ISI_1_) in a 10-spike train. We found that HRs fired fewer spikes per 100 pA current injected compared to LRs (HRs: mean ± SD 8.4 ± 2.6, LRs: 11.7 ± 4.6; Mann-Whitney U = 45, p = 0.0166), indicating reduced excitability in this population. Other measures of excitability were, however, similar for the two groups (Fig. 6).

**Fig. 6.**
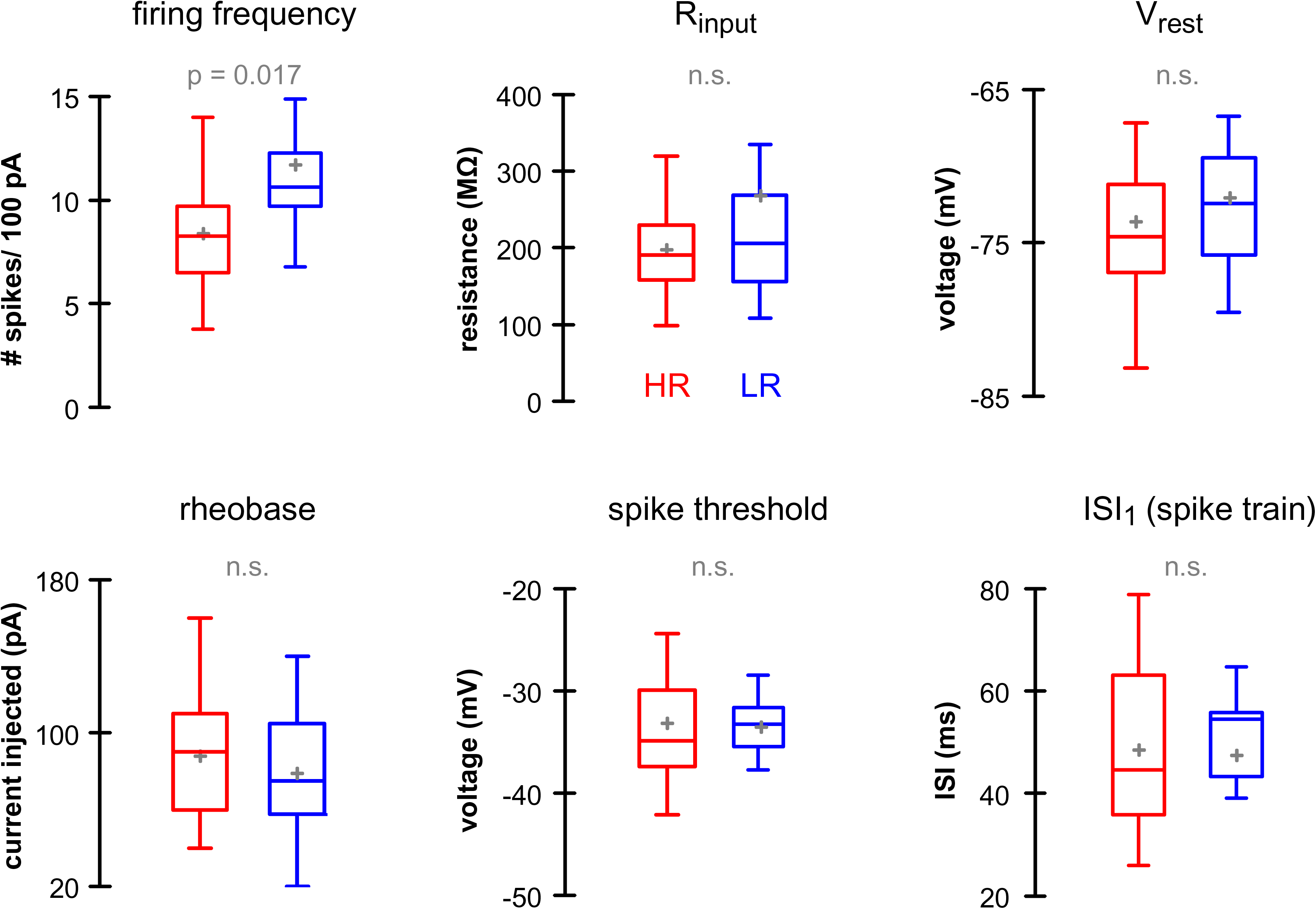
Electrophysiological parameters of neuronal excitability. HR neurons (red; n = 16 cells, 12 mice) and LR neurons (blue; n = 12 cells, 9 mice) were compared in terms of firing frequency per 100 pA injected, input resistance (R_input_), resting membrane potential (V_rest_), rheobase current, spike threshold, and first inter-interspike interval (ISI_1_ in a 10-spike train). Statistical comparisons were performed using a Mann-Whitney *U* test. Box plots show the median (central horizontal bar), mean (+), as well as the first and third quartiles (lower and upper limits).

To further characterize HRs and LRs, we measured passive and active electrophysiological properties. Since excitatory neurons in barrel cortex L2/3 show similar electrophysiological properties (Avermann et al. 2012), we did not expect to see differences between the groups of excitatory HRs and LRs. Indeed, passive membrane properties were comparable in both groups, as demonstrated by similar membrane time constants and sag amplitudes (Table 1). Moreover, no differences were found in action potential (AP) amplitude or half-width, and in the amplitude and timing of the afterhyperpolarization (AHP). Other active properties, including the maximum firing frequency and adaptation of the inter-spike intervals were also comparable in HRs and LRs (Table 1).

**Table 1.**
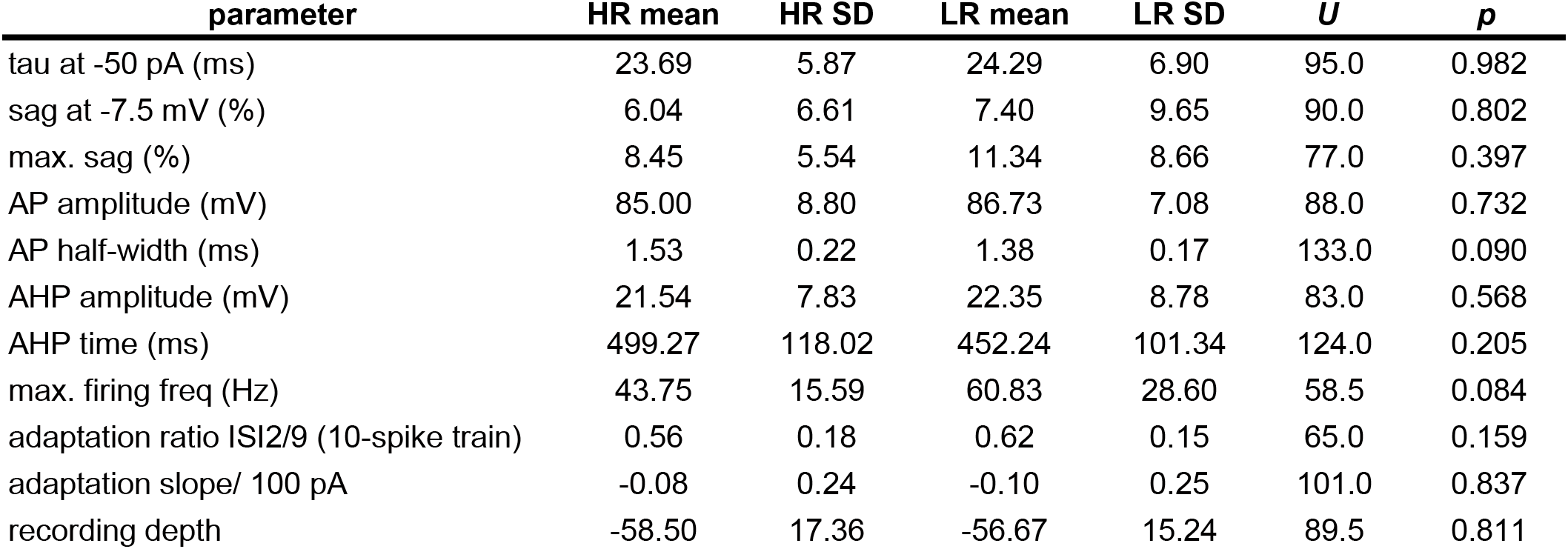
Electrophysiological parameters in tagged HR and LR pyramidal neurons. Comparison between HRs (n = 16) and LRs (n = 12) were assessed using Mann-Whitney *U* tests.

### High responders receive larger and more frequent excitatory network inputs

It is possible that decreased excitability in HRs is a result of homeostatic adaptation to increased inputs (Turrigiano 2012) provided by the network of highly connected HRs. We therefore examined the timing and magnitude of excitatory postsynaptic currents (EPSCs) in high and low responding neurons. Cells were clamped at −70 mV to minimize contamination by inhibitory postsynaptic potentials. Importantly, blockers of spiking activity were not added to the extracellular solution to maintain spontaneous network activity. In total, we recorded 700 s of spontaneous activity in HRs (n = 8 neurons; 5 mice; 3092 EPSCs; see example in Fig. 7A) and 720 s in LRs (n = 8 neurons; 6 mice; 3145 EPSCs; see example in Fig. 7B). For each group, we pooled EPSCs across neurons and found a significant increase in EPSC amplitude in HRs (mean ± SD, −9.17 ± 5.87 pA, n = 3092) compared to LRs (−8.21 ± 4.16 pA, n = 3145; U = 3433278, p < 0.0001; Fig. 7C,D). The increase in HR EPSC amplitude cannot be explained by changes in EPSC properties as the 20-80 % rise time was comparable across groups (HRs: 0.82 ± 0.54 ms, n = 3092; LRs: 0.8 ± 0.53 ms, 3145; U = 4958037, p > 0.05). In addition, HRs showed a slightly reduced inter-event interval of EPSCs compared to LRs (HR: 225.86 ± 239.02 ms, n = 3084; LRs: 228.98 ± 281.8 ms, n = 3137; U = 5035278.5, p = 0.005; Fig. 7C,D), implying more frequent excitatory inputs in HRs compared to LRs. The finding is in line with a slightly higher rate of EPSCs in HRs (4.42 Hz) compared to LRs (4.37 Hz) when looking at the total recording period in both groups. Notably, high-amplitude EPSCs (< −30 pA) were more common in HRs than LRs (Fig. 7C, left panel, inset). These rare, high-amplitude events may play a decisive role in determining overall network activity and may thus be of key importance in stimulus coding (Lefort et al. 2009). Overall, the observed increase in spontaneous excitatory inputs in HRs suggests a presynaptic mechanism for the increased activity level of HRs and lends further support to the idea that HRs form a network of highly active and strongly interconnected neurons.

**Fig. 7.**
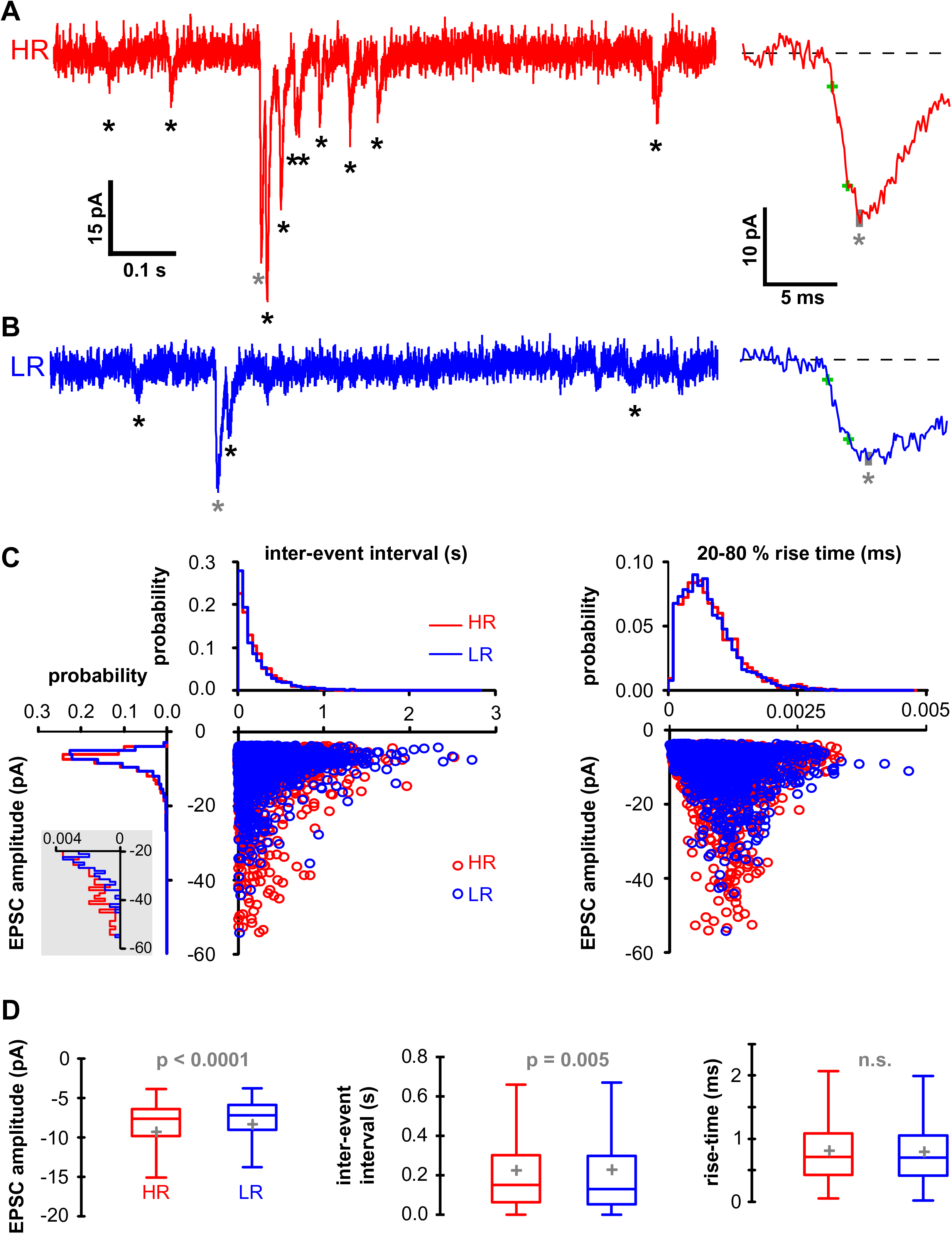
Spontaneous excitatory postsynaptic currents (EPSCs) in HRs and LRs. **(A)** *Left panel:* HR example raw trace with EPSCs (market by asterisks). *Right panel*: Enlarged view of example EPSC (marked by grey asterisk in raw trace) with the 20-80 % rise time indicated by green crosses and the peak indicated by a grey bar. **(B)** LR example raw trace (*left*) and enlarged EPSC (*right*). Same conventions as in *A*. **(C)** Scattergrams of EPSC amplitudes as a function of inter-event intervals (*left*) and rise times (*right*) with respective marginal histograms (50 bins). **(D)** Comparison of amplitude (*left*), inter-event interval (*middle*) and 20-80 % rise time (*right*) between HR EPSCs (n = 3092; 8 cells in 5 mice) and LR EPSCs (n = 3145; 8 cells in 6 mice) using Mann-Whitney *U* tests. Same conventions for box plots as in Fig. 6.

### Morphological characteristics are similar in high and low responders

To pinpoint possible structural differences between high and low responsive cells in barrel cortex L2/3, we filled patched HRs and LRs with biocytin and reconstructed their morphology in 3D subsequently (see **Methods**). In total, morphological features were obtained from 19 tagged excitatory HRs (n = 16 mice) and 10 tagged excitatory LRs (n = 7 mice). In agreement with earlier studies (Feldmeyer 2012; Feldmeyer et al. 2006; Oberlaender et al. 2012), neurons in both groups could be divided in three sub-categories of L2/3 excitatory neurons based on the structure of the apical dendrite: horizontal, broad-tufted and thin-tufted pyramidal neurons (Fig. 8A-C). Horizontal pyramidal neurons were observed in upper L2/3 (mean ± SD, 128.7 ± 20.0 µm below pia; n = 9) and were characterized by an apical dendrite with an atypical, oblique orientation relative to the pial surface. The remaining neurons were classified based on morphological characteristics of the apical dendrites: Neurons for which the field span of the apical dendrites was larger than that of the basal dendrites (see **Methods**) were termed “broad-tufted”, whereas neurons with the inverse relationship were termed “thin-tufted”. In line with previous reports (Feldmeyer 2012; Feldmeyer et al. 2006; Lubke et al. 2003; Oberlaender et al. 2012), broad-tufted neurons were mainly located at an intermediate depth of L2/3 (166 ± 37.8 µm below pia; n = 13), and thin-tufted neurons were more frequent in deeper regions of L2/3 (186 ± 23.5 µm below pia; n = 7; Fig. 8D).

**Fig. 8.**
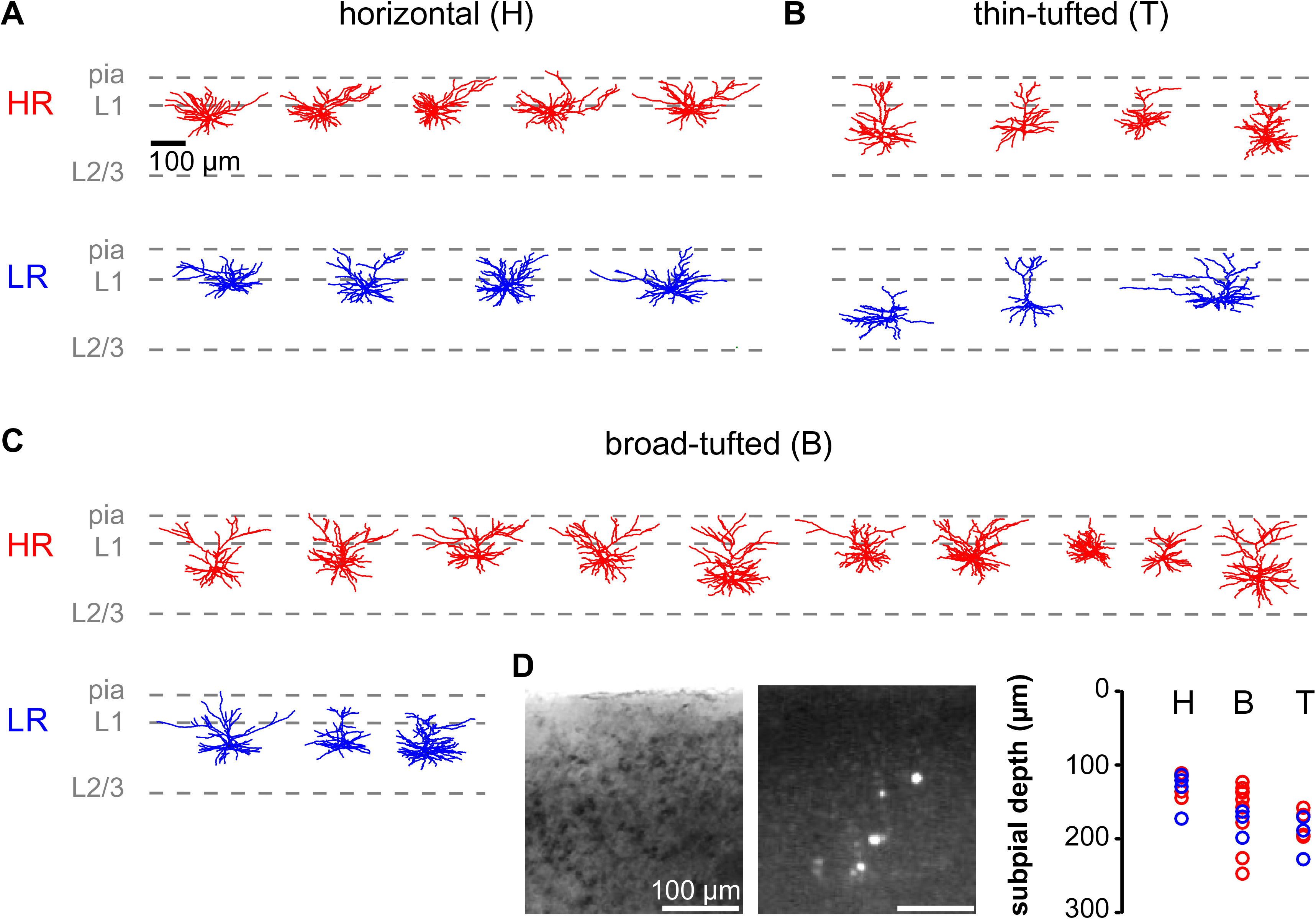
Dendritic morphology of excitatory HRs and LRs. Pyramidal HR neurons (red; n = 19 cells, 16 mice) and LR neurons (blue; n = 10 cells, 7 mice) could be subdivided into three categories based on the structure of the apical dendrite: horizontal, thin-tufted and broad-tufted. **(A)** Pyramidal neurons with a horizontally oriented apical dendrite. Layer borders are indicated by gray dashed lines. **(B)** Pyramidal neurons with a thin-tufted apical dendrite. **(C)** Pyramidal neurons with a thick-tufted apical dendrite. **(D)** Subpial depth of reconstructed cells. *Left-to-right*: Infrared-DIC micrograph of layers 1-3, fluorescent image of tagged cells in layers 1-3, and depth distribution of cells included in morphological analysis (H: horizontal, B: broad-tufted, T: thin-tufted apical dendrites).

Various morphological parameters were examined and compared between HRs and LRs, including the number, length, layer distribution, complexity and spine density of apical and basal dendrites. In addition, the dendritic branching frequency as well as the ratio of apical and basal dendritic length was analyzed. Since no significant differences were found when comparing individual sub-types of excitatory HRs and LRs, the data were pooled and only comparisons at the group level are shown here. However, none of the analyzed morphological parameters differed significantly between excitatory HRs and LRs (Table 2). The recording depth relative to the pial surface was comparable between the two groups, as the patched neurons were located at similar depths in L2/3 (Table 2, Fig. 8D). Our results indicate that morphological characteristics are unlikely to account for the observed differences in activity in HRs and LRs.

**Table 2.** Morphological parameters in tagged HR and LR pyramidal neurons. Statistical comparisons were performed with Mann-Whitney *U* tests.

|  | parameter | HR n | HR mean | HR SD | LR n | LR mean | LR SD | *U* | *p* |
| --- | --- | --- | --- | --- | --- | --- | --- | --- | --- |
| basal dendrites | number | 19 | 7.32 | 1.60 | 10 | 6.80 | 1.40 | 114.5 | 0.372 |
|  | total length | 19 | 2601.43 | 635.50 | 10 | 2519.06 | 1010.16 | 106.0 | 0.636 |
|  | L1 length (% total) | 19 | 12.22 | 20.40 | 10 | 11.14 | 14.39 | 92.0 | 0.903 |
|  | L2/3 length (% total) | 19 | 84.69 | 21.33 | 10 | 84.40 | 14.11 | 106.0 | 0.626 |
|  | complexity | 19 | 37288.37 | 15363.20 | 10 | 49369.52 | 40354.79 | 85.0 | 0.668 |
| | spines/ 100 μm | 14 | 51.43 | 14.50 | 9 | 62.37 | 16.13 | 38.0 | 0.124 |
| apical dendrite | total length | 19 | 2077.47 | 794.02 | 10 | 1944.24 | 612.07 | 105.0 | 0.668 |
|  | L1 length (% total) | 19 | 51.94 | 26.01 | 10 | 52.32 | 31.22 | 94.0 | 0.982 |
|  | L2/3 length (% total) | 19 | 47.77 | 26.38 | 10 | 46.34 | 32.22 | 99.0 | 0.875 |
|  | complexity | 19 | 334794.95 | 249281.69 | 10 | 312926.49 | 242597.45 | 97.0 | 0.946 |
| | spines/ 100 μm | 14 | 55.28 | 17.59 | 9 | 61.78 | 15.63 | 44.0 | 0.250 |
| basal + apical dendrites | branching frequency (nodes/ 100 μm) | 19 | 1.02 | 0.44 | 10 | 0.96 | 0.13 | 72.0 | 0.308 |
|  | ratio total dendritic length (apical/ basal) | 19 | 0.84 | 0.32 | 10 | 0.88 | 0.35 | 87.0 | 0.735 |
| cell body | soma-pia distance (μm) | 19 | 156.25 | 37.76 | 10 | 165.48 | 35.96 | 77.0 | 0.530 |

## Discussion

Our study sheds light on how L2/3 neurons are recruited into active spiking ensembles during passive whisker touch and establishes for the first time a direct link between highly active and less active neurons observed *in vivo* and their characteristics studied *in vitro*. In summary, we found that a small subset (5.7 %) of highly active neurons accounted for a large portion (43 %) of the overall spiking activity measured during passive whisker stimulation. These high responders (HRs) displayed large, fast and reliable activity following stimulation. HRs tended to be co-activated and were more strongly coupled to the population response compared to low responders (LRs). *In vitro* recordings of HRs previously identified *in vivo* revealed an increased amplitude and frequency of EPSCs compared to LRs. The increased excitatory drive in HRs was accompanied by a reduction in intrinsic excitability, as measured by a reduced number of spikes in response to current injection. Both HRs and LRs displayed typical excitatory morphology, including horizontal, thin- and broad-tufted apical dendrites.

In response to principal whisker deflection, excitatory neurons in barrel cortex normally fire only very few action potentials (see e.g. Brecht et al. 2003; de Kock et al. 2007; Kerr et al. 2007; Sato et al. 2007; Simons 1978; Wilent and Contreras 2005), even when optimal stimulation parameters are chosen (Ramirez et al. 2014). This form of sparseness has been referred to as ‘population sparseness’ and is related to a high trial-to-trial variability in spiking output for a given neuron. Another type of sparseness is ‘life-time sparseness’ where a small and fixed subset of neurons responds faithfully to stimulation (Barth and Poulet 2012; Sachdev et al. 2012; Wolfe et al. 2010). This type of sparseness is especially prevalent in barrel cortex L2/3 where studies have identified a neuronal subpopulation that shows elevated stimulus-evoked and spontaneous responses (Kerr et al. 2007; Margolis et al. 2012; Mayrhofer et al. 2015b). Our data provide a link between the two concepts of sparseness: Compared to LRs, HRs show larger and more reliable responses (Fig. 2E,F and Fig. 3A), and contribute more to the stimulus-evoked activity despite fluctuations in the responding ensemble size (*i.e*. trial-to-trial variability; Fig. 3E and Fig. 4B). Thus, lifetime and population sparseness are not mutually exclusive and both can be observed in barrel cortex L2/3.

Apart from being highly responsive to stimuli, HRs also showed more correlated evoked responses compared to LRs (Fig. 4D-F), suggesting that they tended to be co-activated during stimulation. Thus, not only were HRs active during more trials, but they tended to be active during the *same* trials. This shared variability in trial-to-trial activation could originate from either direct synaptic connections or from common inputs (Cohen and Kohn 2011; Shadlen and Newsome 1998). The observation that HR activity is more correlated not only during stimulation but also outside periods of stimulation (Fig. 4G-J) indicates that shared connections, rather than stimulus-driven bottom-up input, may be responsible for their co-activation. In line with this reasoning, correlated spontaneous activity in barrel cortex has previously been shown to emerge from synchronized intracortical inputs, rather than from thalamic inputs (Cohen-Kashi Malina et al. 2016). Furthermore, HRs displayed increased coupling to the population activity (Fig. 4K), which has previously been associated with an increased amount of non-specific excitatory connections with neighboring neurons (Okun et al. 2015). Population coupling predicted a substantial amount of the observed spontaneous correlations in all types of responders (Fig. 4N,O), but left positively-tailed correlation residuals only in HR pairs (Fig. 4P). In V1, such residuals were shown to be mainly dependent on the similarity of the neurons’ tuning curves (Okun et al. 2015). Most importantly, it has been shown that enhanced pair-wise correlations, especially in neurons with similar tuning, directly relate to an increased synaptic connection strength (Cossell et al. 2015; Ko et al. 2011). Taken together, these findings suggest that HRs form a subnetwork of neurons that are highly interconnected.

Increased spontaneous activity and correlated spontaneous events in HRs indicate that their intrinsic or morphological properties might be key determinants for their recruitment during stimulation. Previous studies addressing this issue in barrel cortex L2/3 identified HRs *in vitro* based on elevated expression of the activity-marker *c-fos* (Benedetti et al. 2013; Yassin et al. 2010) or based on electrical stimulation of L4 (Elstrott et al. 2014). In agreement with our results (Fig. 6), these studies found a reduction in intrinsic excitability in HRs as evidenced by a smaller slope of the IF curve, while resting membrane potential and input resistance did not differ from LRs (Elstrott et al. 2014; Yassin et al. 2010). However, the studies disagreed on potential differences in rheobase current and action potential threshold between HR and LRs. Our study design allowed us to address this issue by identifying HRs *in vivo* and measuring their properties *in vitro*, thereby avoiding possible methodological confounds in previous research. Our results indicate that rheobase current and action potential threshold are unaltered in HRs (Fig. 6).

If HRs are less excitable than LRs, then more inputs may explain their enhanced responsiveness. In line with previous *in vitro* research (Yassin et al. 2010), we found that HRs show a higher frequency of spontaneous EPSCs. In addition, our results also revealed larger amplitudes of EPSCs in HRs (Fig. 7). These findings indicate that HRs may down-regulate their intrinsic excitability in response to elevated excitatory inputs in order to maintain their homeostatic balance (Gainey and Feldman 2017; Lambo and Turrigiano 2013), as previously suggested (Elstrott et al. 2014; Yassin et al. 2010). The highly skewed distribution of EPSC amplitudes, with many small-amplitude and few large-amplitude events, is in close agreement with previous data on excitatory postsynaptic potentials (EPSPs) in barrel cortex L2/3 (Feldmeyer et al. 2006; Holmgren et al. 2003; Lefort et al. 2009). Based on experimental observations and computational models, it has been argued that rare large-amplitude events may be more important than small ones in determining the overall network activity and may be fundamental to sparse stimulus coding (Lefort et al. 2009; Poulet and Petersen 2008). In this context, it is interesting to note that HRs received a higher frequency of large EPSCs compared to LRs (Fig. 7C, left panel, inset), indicating that HRs may function as ‘hubs’ in processing sensory information. The observed enhanced correlations among HRs *in vivo* (Fig. 4D-J) support the notion that large-amplitude EPSCs may arise as a result of correlated activity of pre- and postsynaptic neurons (Feldman 2000; Hebb 1961; Markram et al. 1997; Sjostrom et al. 2001). In contrast, low-amplitude EPSCs may provide opportunities for plasticity in the neural network, as previously suggested (Lefort et al. 2009).

An increase in excitatory inputs to HRs may arise from enhanced bottom-up, top-down or intracortical drive. In our preparation of thalamocortical brain slices, inputs from longer-range connections (even outside S1) cannot be completely excluded but are rather unlikely due to the truncation of axons during the slicing process. It is more likely that the increased inputs to HRs arose from local L4-to-L2/3 or L2/3-to-L2/3 connections because of the high density of presynaptic L4 and L2/3 axons and postsynaptic L3 dendrites, which results in a large number of potential synaptic connections (Brecht et al. 2003; Feldmeyer et al. 2006; Feldmeyer et al. 2002; Holmgren et al. 2003). Previous studies discovered that c-fos-labeled HRs in L2/3 were more likely to be connected to each other (Yassin et al. 2010) and received a larger number of L4 inputs compared to LRs (Benedetti et al. 2013). Similarly, studies in visual cortex revealed that if L2/3 pyramidal cells shared common input from L4 and within L2/3, they were more likely to be reciprocally connected (Yoshimura et al. 2005). These findings support the idea that enhanced inputs to HRs arise from L4 as well as L2/3, in particular from other connected L2/3 HRs.

As differences in neuronal function may also relate to differences in morphological features (Oberlaender et al. 2012), we set out to characterize the neuronal structure of HR and LRs by means of biocytin staining and 3D morphological reconstructions. Our study showed for the first time that HRs span all three subcategories of pyramidal cell morphologies, including thick-tufted, thin-tufted and horizontal pyramidal neurons (Fig. 8). Similar to an earlier study (Elstrott et al. 2014), HRs and LRs did not show differences in the spine density or total length of basal or apical dendrites. In addition, we found no differences in the complexity, layer-length distribution and branching frequency of basal and apical dendrites. The similar morphology of HR and LR pyramidal neurons indicates that differences in responsiveness may be largely due to functional characteristics related to connectivity and enhanced inputs.

An often-debated issue is whether the observed sparseness of responses is a result of anesthesia, during which activity in the brain is generally suppressed (Barth and Poulet 2012). We believe that anesthesia-related effects did not play a significant role in our study since the concentration of administered isoflurane was kept to a minimum in order to observe large stimulus-evoked responses. Moreover, several studies have previously demonstrated sparse activity of L2/3 excitatory cells both during anesthetized and awake states (Crochet et al. 2011; de Kock and Sakmann 2009; Margolis et al. 2012; O’Connor et al. 2010), indicating that sparse responses are a general feature of L2/3 cells and not exclusively associated with the anesthetized brain state.

Sparse population responses are not unique to the barrel cortex, but occur across different brain regions and in different species (Hromadka et al. 2008; Ko et al. 2011; Okun et al. 2015; Poo and Isaacson 2009; Vinje and Gallant 2000). It has been argued that the advantage of sparse coding is the reduced metabolic cost associated with a reduced number of firing neurons (Laughlin 2001; Laughlin and Sejnowski 2003; Lennie 2003). However, it has been shown that AP generation is not as metabolically costly as has been previously thought (Alle et al. 2009). In addition, maintaining a large number of non-responsive neurons in a resting state is also energy-expensive. Thus, there might be other, more important benefits associated with a sparse code, *e.g*. that it allows for transmitting signals with a higher information content (Olshausen and Field 2004) or that it represents a salient signal in the brain (Wolfe et al. 2010). Indeed, the firing of a few L2/3 barrel cortex neurons has been shown to be behaviorally detectable by the animal (O’Connor et al. 2010). In the future, it will be interesting to determine the effects of activating or de-activating HRs under different behavioral conditions, *e.g*. when the animal is solving a discrimination task, and to track HR activity under the influence of plasticity, *e.g*. after whisker removal. Optogenetic tools in combination with two-photon holographic stimulation (Chaigneau et al. 2016) can help to shed light on these questions.

## Acknowledgements

The RCaMP2 construct has been kindly provided Haruhiko Bito (University of Tokyo, Graduate School of Medicine, Tokyo, Japan). pAAV.Syn.NES-jRGECO1a.WPRE.SV40 was a gift from Douglas Kim & GENIE Project and pACAGW-H2B-PAGFP-AAV was a gift from Massimo Scanziani. We thank Werner Hucko (Research Centre Jülich, Jülich, Germany) for the histological processing of the brain slices and Axel Honné (Medical School, RWTH Aachen, Aachen, Germany) for technical support. Furthermore, we thank Björn Michael Kampa (RWTH Aachen University, Aachen, Germany), Charly Rousseau and Christoph Schmidt-Hieber (Institut Pasteur, Paris, France) for useful discussions on the manuscript. This work was supported by the Minerva Stiftung Gesellschaft für die Forschung mbH (D.G.), SFB-1213 Project B01 (S.R.), the Helmholtz Society (D.F.), the DFG Research Unit ‘‘Barrel Cortex Function” (Grant 472/4-2; D.F.), the EU’s Horizon 2020 Research and Innovation Programme (Grant Agreement No. 720270, HBP SGA1; D.F.), and the IZKF Aachen, Medical Faculty of RWTH Aachen University, Germany (F.H.).

## Author contributions

Conceptualization, F.H.; Methodology, C.B., P.M.G. and F.H.; Software, C.B., P.M.G. and D.G.; Formal Analysis, C.B. and P.M.G.; Investigation, C.B. and P.M.G; Resources, D.G., S.R., D.F. and F.H.; Writing – Original Draft, C.B.; Writing – Review & Editing, C.B., P.M.G., D.G., S.R., D.F. and F.H.; Visualization, C.B. and P.M.G.; Supervision, D.F. and F.H.; Project Administration, C.B.; Funding Acquisition, D.F. and F.H.

## Data availability

The data that support the findings of this study are available from the corresponding author upon reasonable request.

## Methods

### Animals

Animal procedures were approved by the local authorities (animal facility at University Hospital Aachen and the Northrhine-Westphalian Landesamt für Natur, Umwelt und Verbraucherschutz) and were carried out in accordance with the German Animal Welfare Act, the European Directive on the Protection of Animals used for Scientific Purposes, as well as guidelines by the Federation of European Laboratory Animal Science Associations (FELASA). Thirty C57/BL6J adult, male mice were used for the procedures. Mice were obtained from Charles River and kept under a 12h light-dark cycle, with food and water available *ad libitum*.

### Surgery

At the time of surgery, animals were aged between P45 and P105. Mice were anesthetized with isoflurane (AbbVie, Germany; induction: 4 %, maintenance: 2 %) and placed in a stereotactic frame (Kopf Instruments, USA). They were injected with buprenorphine (Temgesic; Reckitt Benckiser, UK; 0.02 mg/kg, s.c.) for pain management and their eyes were protected from desiccation by applying ointment. The fur was removed over the skull, the skin was disinfected with betadine (Mundipharma, Germany), and bupivacaine (Actavis, Germany; 0.25%, 0.04 ml) was injected subcutaneously for local analgesia. The skull was exposed and the C1 barrel in the left hemisphere was identified using intrinsic optical imaging and piezo-controlled whisker stimulation (Margolis et al. 2012; Mayrhofer et al. 2015). A 3 x 3 mm craniotomy was performed and two viruses were injected sequentially in L2/3 of the C1 and adjacent barrel columns (Fig. 1,2): One virus for expressing histone-associated, photo-activatable green fluorescent protein (pACAGW-H2B-PAGFP-AAV was a gift from Massimo Scanziani [Addgene plasmid # 33000; http://n2t.net/addgene:33000; RRID:Addgene_33000]; AAV9 was produced by Viral Core Facility Charite Berlin, Germany) that was used for photo-tagging cells (Lien and Scanziani 2011), and a red calcium indicator for two-photon imaging of neuronal activity (Dana et al. 2016; Inoue et al. 2015). The red indicator was either jRGECO1a (23 animals; AAV1.Syn.NES.jRGECO1a.WPRE.SV40; Dana et al. 2016) obtained from Penn Vector Core, USA, or RCaMP2 (5 animals; AAV-hsyn-RCaMP2) obtained from the Viral Core Facility of the Charite Berlin, Germany (see below). Virus stocks showed high titers (PAGFP: 5.36e12 GC/ml; jRGeco1a: 2.08e13 GC/ml; RCaMP2: 5,01e12 VG/ml) and were diluted 1:1 with mannitol (Mannit 15 %; Serag, Germany) for injections. To avoid tissue damage, small volumes of virus were injected (PAGFP: 400-450 nl; jRGeco1a: 600-700 nl; RGeco: 600-650 nl) at slow rates (∼ 50 nl/min). Injections were performed using beveled glass pipettes (Drummond, USA) with a diameter of 13-20 µm. The craniotomy was then covered with a glass window (3 x 3 mm; UQG Optics Ltd, UK). Dental cement (DE Healthcare Products, UK) was applied to keep the window in place and to form a head cap holding a custom-made head-post made of aluminum or titanium. Throughout surgery, body temperature was maintained at 37 °C with a feedback-controlled heating pad (Thorlabs, USA). After surgery, mice received buprenorphine injections for pain management (Temgesic; Reckitt Benckiser, UK; 0.02 mg/kg s.c. every 12 h, 3 d postoperatively). Antibiotic treatment was administered one day before surgery until the end of the experiment by supplementing the drinking with enrofloxacin (Baytril, 1 ml/l; Bayer, Germany). Post-operative pain management was carried out using buprenorphine (Temgesic; Reckitt Benckiser, UK; 0.02 mg/kg, s.c.; 2 x per day for 1-1.5 days). Mice were single-housed for the rest of the experimental procedures to avoid potential damage to the implant.

### AAV-hsyn-RCaMP2 virus generation

A plasmid containing the coding sequence (cds) of *R-CaMP2* was kindly provided by Haruhiko Bito (Inoue et al. 2015). Using PCR, the cds was complemented with a BamHI and Kozak sequence upstream of the translational start site and an AscI restriction site behind the stop codon. The resulting product was subsequently used to replace the cds of *G-CaMP6-p2a-nls-dTomato* in plasmid #51084 obtained from *Addgene*. The resulting vector allows the cell type specific expression of R-CaMP2 under the human *synapsin 1* regulatory element and delivery through AAVs. Primer sequences used for amplification of *R-CaMP2* were:

Fwd: 5’-GGATCCGCCACCATGGGCTCTCACCATCACCATCACCATGGAATGGCTAGCATGACTGGTGGA-3’;

Rev: 5’-GGCGCGCCCTAGCTGCTGATGGCGTGTCAGAACTATGGGTTGGACTCCAC GTCTCC-3’

### In vivo two-photon imaging

#### Intrinsic optical imaging

Prior to two-photon calcium imaging, intrinsic optical imaging (IOI) was performed to identify the single whisker representation in neocortex using a 50 mm tandem lens system. Imaging was conducted at 30 Hz with a 12-bit CCD camera such that the field of view (FOV) spanned the entire cortical region covered by the window. The imaged region was continuously illuminated with a red-light emitting diode (630 nm wavelength). For stimulation, a single whisker was inserted into a glass capillary mounted on a piezoelectric element powered by a piezo-controller (MTD693B, Thorlabs, USA). The whisker was moved rostro-caudally for 6 s with a 10 Hz square wave pulse (amplitude: ∼ 0.5 mm). This stimulation protocol was repeated at least 10 times with inter-trial intervals (ITI) of 20 s. The change in absorbed light was computed as

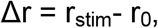

with r_0_ being the average image in the 5 s window prior to stimulation and r_stim_ the average image in the 5 s window prior to the end of stimulation. Δr revealed discrete areas of functional activity corresponding to the single whisker representation (Fig. 2A). A green-light emitting diode (530 nm) was used to visualize the vasculature pattern in order to match IOI images to subsequent two-photon imaging.

#### Two-photon calcium imaging data acquisition

Two-photon imaging was carried out after a minimum of two weeks following surgery to allow for sufficient viral expression. Functional images were acquired continuously using bidirectional galvanometer scanning mounted in a custom built two-photon microscope (Mayrhofer et al. 2015a). The imaging rate was set to either 8.5 Hz or 11.4 Hz with an image resolution of 256*256 or 256*220 pixels, respectively, spanning ∼ 200 μm^2^. The FOV was placed on the center of the IOI signal. The genetically encoded red calcium indicator (jRGECO1a or RCaMP2) was excited with a Ti:sapphire laser (1040 nm, Chameleon Ultra II; Coherent, USA) using 25-60 mW depending on the recording depth. Emitted light was recorded through a 16x, 0.8 NA objective (Nikon, Japan) and detected with two Hamamatsu photomultiplier modules (H10770PA-40) with the following filter settings: band pass filter 540/40 nm and 617/73 nm for green and red channels, respectively, and a 750 nm short pass filter (Semrock, USA) for each channel. Two-photon laser scanning and image acquisition was performed using version r3.8 or 2017 of the ScanImage software (Vidrio technology, USA, Pologruto et al. 2003).

Animals remained anesthetized during the whole procedure with isoflurane (AbbVie, Germany; induction: 4 %, maintenance: ∼ 0.5-1.5 %) and head-fixed. Body temperature was maintained at 37 °C with a feedback-controlled heating-pad and respiratory rate was monitored by means of a piezoelectric sensor placed under the animal’s trunk and a custom-written LabView program (National Instrument, USA). Images were acquired when the respiratory rate was above 120 breaths per minute and whiskers were still, as assessed with a camera placed underneath the animal’s head. One target whisker at a time (B1, C1, D1, C2 or D2) was placed into a capillary glass tube that was mounted on a piezo element and positioned 3 mm away from the whisker pad. Whiskers other than the stimulated one were either trimmed below 3 mm length or were located at sufficient distance to not be in contact with the whisker stimulation apparatus. The stimulus consisted of a periodic rostro-caudal deflection of the target whisker with a prototypic pulse (120 Hz sine wave, 700 °.s^-1^, ∼ 0.5 mm amplitude) that was delivered at 90 Hz for 1 s, as previously described (Mayrhofer et al. 2015b). This was repeated 60 to 100 times with an ITI of 6.2 s. Multiple FOVs could be acquired sequentially in the same animal, either regularly spaced in the z-dimension (1-5 different depths, at 100 to 300 μm from the pial surface), and/or in several whiskers barrel fields. In a subset of 21 FOVs from 6 animals with jRGECO1a, extra epochs of spontaneous neuronal activity lasting between 175 s and 1000 s were recorded immediately following the stimulation session. No sensory stimulation was delivered to the whiskers during these epochs. Immediately after imaging a FOV, the calcium imaging data was analyzed, and target cells were selected and photo-converted.

#### Two-photon calcium imaging data processing

Movies were first motion-corrected using the registration module from suite2p (Pachitariu et al. 2016). We then manually selected region of interests (ROIs) corresponding to individual neurons using a custom Matlab software (Mathworks, USA). Selection and segmentation was based on three average frames: The first frame was an average of all frames in the red channel to see expression of the calcium indicator. The second frame was an average of all frames in the green channel to visualize nuclei trough PAGFP expression. The third frame was a Δf/f_0_ heat map based on the pixel-wise response to the stimulation in the red channel, which was computed as

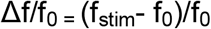

where f_0_ was the average pixel value in frames during the baseline (1 s prior to stimulus start), and f_stim_ was the average pixel value in frames during the 1 s stimulation.

Immediately after calcium imaging acquisition in a FOV, only cells that met the criteria for photo-conversion were selected. For detailed analyses of calcium imaging data presented in Fig. 2-4 (carried on the subset of 21 FOV with extra epochs of spontaneous activity), ROIs were selected on every “visible neuron”. A visible neuron was defined as a distinguishable ring of red calcium indicator surrounding a nucleus of green PAGFP, based on averaged green and red frames over the session. Neurons’ individual fluorescence time series were obtained by averaging pixel values in the corresponding ROI. Local, out-of-focus fluorescence (neuropil) was measured by averaging pixel fluorescence values in a ring of ∼ 40 µm diameter surrounding the ROI, excluding pixels from other ROIs (Peron et al. 2015). Slow baseline fluctuations were removed individually from each neuronal and neuropil time series by subtracting the 10th percentile in a 30 s wide moving window. Neuropil subtraction was then scaled for each individual ROI to the slope *α* according to the following equation:

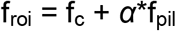

where f_roi_ represents the variation of fluorescence observed in the ROI, f_c_ denotes fluorescence originating from the cellular soma, *α* is the scaling factor of neuropil contamination and f_pil_ is the neuropil fluorescence. Most values of *α* were found in a range of 0.45-0.9 with a robust linear regression (Driscoll et al. 2017). Values below and above this limit were set to a minimum of 0.45 and a maximum of 0.9, respectively. We used this corrected fluorescence trace as a common basis for the computation of Δf/f_0_ time series, z-scored time series and spiking estimates. The Δf/f_0_ time series was computed individually for each trial as

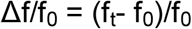

where f_0_ was the average fluorescence value during the baseline (1 s prior to stimulus start) and f_t_ was the average fluorescence in frame t. The peak of Δf/f_0_ was the maximal value within the stimulation period after averaging Δf/f_0_ across trials. Z-score time series were computed as

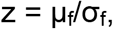

with σ_f_ = f_16_ - f_2.3_

and µ_f_ = f_2.3_ +2* σ_f_

where f_16_ is the 16^th^ percentile of all fluorescence values in the time series and f_2.3_ is the 2.3^th^ percentile. These values were picked because the distance between them represents roughly 1 SD in Gaussian distributions. Gaussian noise statistics were computed on the negative hillside of the fluorescence histogram (representing the non-active period of the neuron), such that z-score time series were normalized to the noise level independent of the cellular activity level. Therefore, z-scores could be used to detect calcium event as follows: When two consecutive time points had a z-score above 2.5, it was considered as a calcium event. Rather than retrieving precise spike timing, this analysis focused on estimating key statistical parameters to reliably detect periods of activity. These calcium events were later used in binary form over 1 s time bins to study neuronal co-activation in the recorded population ensemble. Finally, a constrained deconvolution was applied on the corrected fluorescence trace (Pachitariu et al. 2016) to obtain a continuous spiking estimate. This spiking estimate was used for later computation of pair-wise correlations. These methods and criteria were optimized from the freely available jRGECO1a dataset from the CRCNS website (Mohar et al., 2016; Dana et al. 2016).

#### Target cell selection

Neurons were classified as high responders (HRs) when they met three criteria: First, their average Δf/f_0_ peak across trials had to cross a threshold of 25 % during the stimulation period. Second, these neurons had to show significant activation in at least 25 % of the trials. This was assessed by a two sample t-test of the activity during baseline (1 s before stimulation) versus the fluorescence during 1 s stimulation. Third, a significant increase in fluorescence had to occur in the 2 imaging frames following the onset of stimulation (at either sampling rate). This was tested across active trials with a 2-sample t-test between Δf/f_0_ in the n^th^ frame versus baseline average. Neurons were classified as low responders (LRs) if their average Δf/f_0_ peak was below a threshold of 10 % and if they showed significant activation in less than 10 % of the trials. Neurons that did not meet criteria for either LRs or HRs were classified as ‘medium responders’ (MRs).

#### Photo-conversion of the PAGFPP

Once the target ROIs were selected, the corresponding PAGFPP expressing cell nuclei were identified at 850 nm excitation wavelength (imaging power ranging from 4-10 mW depending on the recording depth). The precise position of the focal plane in the z-axis was adjusted with μm precision using a piezo actuator controlled through ScanImage. This strategy was needed to compensate a shift in the focal depth of ∼ 10 μm induced by the change in excitation wavelength from 1040 nm to 850 nm. Using a ScanImage feature for sub-FOV imaging, we manually drew rectangles on individual target nuclei with a resolution of 16 x 16 pixels (∼ 4 μm^2^). Target ROIs of 16 x 16 pixels (∼ 4 μm^2^) with a line period of 512 ns were illuminated at 750 nm for 2 s in order to photo-convert PAGFPP. The laser power was adjusted ranging from 5-15 mW depending on cell depth. Each cell was individually and sequentially imaged in order to photo-convert the PAGFPP. Photo-conversion efficiency was assessed at either 950 nm (4-12 mW) or 1040 nm (25-60 mW).

#### Population coupling and correlations

Trial-to-trial correlations, also called noise correlations, were calculated using the Pearson correlation coefficient between two neuronal responses to repeated presentations of the same stimulus (Cohen and Kohn 2011). We used spiking estimates of each cells, excised period of baseline, and averaged the spike estimate within 1 s whisker stimulation period, generating one activity vector per cell. All correlations between pairs of cells were calculated from this activity vector after subtraction of the average response to stimulation. To measure spontaneous correlations during ongoing activity, we adopted the same approach: Spike estimates were averaged in 1 s long, non-overlapping time bins over the entire period of spontaneous activity.

Correlations between population rate and single cell activity were assessed in the same FOV. The fraction of the population that was active in a given trial (also termed “ensemble size”) was calculated as the number of neurons detected active (z-score > 2.5 in two consecutive frames within the stimulation period) divided by the total number of neurons in the FOV. For a single FOV, ensemble size from all trials were pooled, sorted in ascending order and distributed in six bins of equal size. By taking the rank of the ensemble size activation (from 1-6, smallest active ensembles to largest active ensembles), it was possible to pool datasets from multiple FOVs.

Population coupling per cell c_i_ could be measured following the formula from Okun et al., 2015:

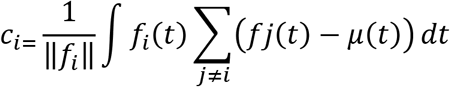

f represents the binary vector of activity, µ its mean event rate and ||f_i_|| its norm (*i.e*. the total sum of events). Couplings were pooled across sessions by z-scoring data separately in every FOV.

Activity shuffling was performed on a binarized activity vector. The independent population activity pattern (Fig. 3A) was generated by randomly shuffling the single neuron activity vector in time, keeping thus the number of events per cell constant, but modifying the trial-to-trial population rate. Generation of activity patterns with preserved mean event rate, population rate, and population coupling was performed as follows (Fig. 4A): From the original matrix of binarized activity, where rows represent neurons and columns represent time bins, every detected event was randomly exchanged once with another random event from a different cell. In other words, rows of 2 x 2 sub-matrices of activity containing two distinct events from two neurons 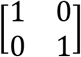 were exchanged - an operation described by Okun et al., 2015. This first step completely randomized the activity matrix while preserving mean event rate and population rate. The third constraint on randomization, (population coupling) was then met using the same approach on 2 x 2 sub-matrices until a convergence threshold was reached. The difference was that the two rows (neurons) were selected to have the opposite deviation of cell coupling from their initial value, one too high and one too low. This had the effect to make the modeled population coupling of each cell to converge towards its initial value (see Okun et al, 2015). The process was stopped when the sum of squared errors from initial coupling was below the number of neurons. Based on the modeled data, we could compute modeled pair-wise correlations in all pairs of cells, and compare them to the experimentally observed pair-wise correlations. To quantify predictions of noise correlations from this model, we used a numerical index D of explained deviation from 0, which was computed as

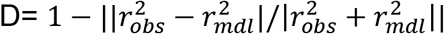

where 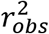 is the experimentally observed Pearson correlation, and 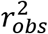 the modeled Pearson correlation. This index D varies between 0 and 1, from incongruent to perfectly congruent prediction from the model, respectively. Every randomized shuffling procedure was repeated at least 250 times.

To avoid spurious relationships between the event rates and correlations when comparing different classes of neurons, we matched geometrical mean event rates (GMFRs) across classes of responders during epochs of spontaneous activity (Fig. 3-4). It was computed as

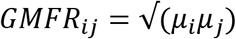

*μ_i_* being the mean spike estimate of neuron i (Fig. 3) or the mean event rate (Fig. 4). Next, we distributed all GMFRs from the three responder’s classes in bins of 0.01 Hz width. From each bin we drew the same number of samples for the three classes. Thus, the numbers of pairs and their distributions of GMFRs were matching across the three responder classes with a resolution of 0.01 Hz.

### Brain slice preparation

After *in vivo* two-photon imaging, mice aged P72 to P213 were anesthetized with isoflurane and decapitated. The brain was quickly removed and placed in 4 °C cold artificial cerebrospinal fluid (ACSF) containing 206 mM sucrose, 2.5 mM KCl, 1.25 mM NaH_2_PO_4_, 3 mM MgCl_2_, 1 mM CaCl_2_, 25 mM NaHCO_3_, 12 mM N-acetyl-L-cysteine and 25 mM glucose (325 mOsm, pH 7,45; aerated with 95 % O_2_ and 5 % CO_2_). Then the brain was shortly placed in a custom-made chamber with a ramp (10 ° in the rostral-caudal direction) to make an initial cut with a razor blade at an angle of 45 ° relative to the midline. The brain was then quickly tranferred to a vibratome (HM 650 V, Thermo Fisher Scientific, Germany) filled with ASCF and cut in 300 µm thick slices. The thickness and slicing angle were optimized for preserving neuronal morphology, which is important for maximizing the number of intact neurons in the slice and for high-quality morphological reconstructions. After cutting, slices were incubated (30 min at 33 °C and then at room temperature) in ACSF containing 125 mM NaCl, 25 mM NaHCO_3_, 2.5 mM KCl, 1.25 mM NaH_2_PO_4_, 5 mM MgCl_2_, 1 mM CaCl_2_, 25 mM glucose, 3 mM Myo-Inositol, 2 mM Na-pyruvate, 12 mM N-acetyl-L-cysteine and 0.4 mM vitamin C (300 mOsm; 95 % O_2_, 5 % CO_2_). This protocol was adapted from previous studies (Ting et al. 2014; van Aerde and Feldmeyer 2015) and modified to obtain healthy slices from adult mice (P30 up to 8 months). Using this approach, we were able to record for about an hour in the slice before the cell quality would start deteriorating.

### In vitro patch clamping

Intracellular recordings were performed at room temperature. This deviation from standard protocols was necessary to maintain the health of the adult slices, presumably because oxygenation is better at lower temperatures. During recordings, slices were perfused at 5 ml/min with an extracellular ACSF containing 125 mM NaCl, 25 mM NaHCO_3_, 2.5 mM KCl, 25 mM glucose, 1.25 mM NaH_2_PO_4_, 2 mM CaCl_2_ and 1 mM MgCl_2_ (300 mOsm; 95 % O_2_, 5 % CO_2_). The intracellular solution contained 135 mM K-gluconate, 4 mM KCl, 10 mM HEPES, 10 mM phosphocreatine-Na, 4 mM ATP-Mg and 0.3 mM GTP (pH 7.3; 290 mOsm). Biocytin (Sigma Aldrich, Germany) was added to the intracellular solution at a concentration of 3 mg/ml and cells were filled during the recordings. Glass pipettes were pulled from borosilicate glass capillaries (2 mm outer diameter, 1 mm inner diameter) with a P-1000 Flaming/Brown micropipette puller (Sutter Instruments, USA) and had a resistance of 5-8 MΩ. Slices were observed with an upright microscope (Axio Examiner D1 with 5x/0.16 plan and 40x/1.0 DIC immersion objectives; Zeiss, Germany) under infrared differential interference contrast (IR-DIC) and fluorescent illumination (BP 500/25, FT 515, BP 535/30; Zeiss, Germany). Signals were acquired with an EPC10 amplifier (HEKA, Germany), sampled at 10 kHz and filtered at 2.9 kHz using Patchmaster software (HEKA, Germany). Offline analysis was carried out using Igor Pro software (Wavemetrics, USA).

### Histology

After patch recordings, brain slices were placed in fixative (4 % paraformaldehyde, 100 mM phosphate-buffered saline; pH 7.4) and stored at 4 °C for a minimum of 48 h. Following a protocol published previously (Marx et al. 2012), biocytin-filled neurons were visualized using avidin-biotinylated horseradish peroxidase complex reaction (ABC-Elite; Camon, Germany) and 3,3-diaminobenzidine (Sigma Aldrich, Germany) as a chromogen. After dehydration and embedding in Eukitt (Marienfeld Laboratory Glassware, Germany), neurons were reconstructed in 3D using Neurolucida® software (MBF Bioscience, USA) under an Olympus Optical BX61 microscope with a final magnification of 1000x.

### Analysis of patch clamp data

Series resistance was measured in whole-cell mode at −60 mV with a −10 mV square pulse applied once a second for 300 ms (10 repetitions), compensated by 80 % and monitored throughout the experiment. Neurons were excluded from the analysis if their series resistance was larger than 60 MΩ or changed by more than 10 % during the experiment. Because the majority of identified high responders were excitatory neurons, only these were included in the analysis. Excitatory neurons were distinguished from GABAergic interneurons based on their firing pattern and morphology. Resting membrane potential (V_rest_) was measured immediately after break-through.

The following passive membrane properties were measured in current-clamp mode by injecting 1 s-long current in steps of 10 pA starting at −100 pA: Input resistance (R_input_) was calculated as the slope of the linear fit of the current-voltage (IV) curve measured close to V_rest_ in a window of −50 to 50 pA. The membrane time constant τ was calculated with a mono-exponential fit of the voltage onset response to a −50 pA current. The voltage sag was defined as the percentage difference between the initial and sustained response to current injection. Two types of voltage sag were acquired: One was measured when the neuron was hyperpolarized by approximately −7.5 mV (“sag at −7.5 mV”), and the other was measured as the maximum voltage excursion independent of the level of hyperpolarization (“max. sag”). The rheobase was determined as the minimum current amplitude required for eliciting an action potential (AP). The first elicited spike was used for quantifying AP properties, including the threshold (second derivative of membrane potential exceeding 3x SD of its baseline value before spike generation), amplitude (difference between peak and threshold membrane potentials), half-width (measured at half-amplitude), and the amplitude and timing of the afterhyperpolarization (AHP; difference between spike threshold and the most negative membrane potential measured during the AHP).

The following active properties were obtained in current-clamp mode by injecting 1 s-long current steps of 25 pA starting at −20 pA: The firing frequency per 100 pA current injected, the maximally measured firing frequency, and the slope of the adaptation measured as the ratio between the 2^nd^ and 9^th^ inter-spike intervals (ISI). In addition, a sweep with a 10-spike train was chosen for every cell to measure doublet spiking (1^st^ ISI) and adaptation (ratio of 2^nd^ and 9^th^ ISIs). Since not all cells fired exactly ten spikes, a range of 9-12 spikes was chosen (if only 9 spikes were available, then the 2^nd^ and the 8^th^ ISI was chosen for calculating adaptation).

Spontaneous excitatory postsynaptic currents (EPSCs) were measured in voltage-clamp at −70 mV holding potential for periods of 70-90 s. Importantly, inhibitors such as tetrodotoxin or gabazine have not been applied in order to maintain spontaneous network activity. In a first step, the mean baseline activity and the standard deviation (SD) of the noise were quantified in a 1 s time window without obvious spontaneous events. Subsequently, a threshold-crossing detection algorithm implemented in the SpAcAn (G. P. Dugué & C. V. Rousseau) extension for Igor Pro was used for detecting EPSCs: For every point *N* of the trace, the algorithm calculated the difference between the baseline segment (8 ms window centered on *N*) and a peak segment (0.3 ms window). Baseline and peak were separated by a rise time segment (0.8 ms window). Whenever the difference exceeded a certain threshold, the algorithm then identified a local maximum in a 3 ms peak detection window. The threshold was set for every cell individually at three times the SD of the noise. All automatically detected peaks were verified by visual inspection. For statistical comparisons, EPSCs from neurons were pooled for each group.

### Analysis of morphological data

Neurons with clear labeling and minimal background staining were chosen for further morphological analysis. Care was taken to include only neurons with minimal to no dendritic truncations as assessed based on the number of dendrites at the slice surface with ‘blebs’ resulting from biocytin spill. Neuronal morphology was reconstructed in 3D using an Olympus BX61 microscope fitted with a 100x oil immersion objective (NA 1.4) and a 10x eye piece. Reconstructions of neurons, pial and layer borders were corrected for tissue shrinkage by a factor of 1.1 in the x- and y-dimension and 2.1 in the z-dimension (Neurolucida software, MBF Bioscience, USA). Thick- and thin-tufted excitatory neurons were distinguished based on the ratio of the apical and basal dendritic field spans. Field spans were measured parallel to the pia as the maximal spread of the dendritic branches. Neurons with apical/dendritic field span ratios > 1 were defined as thick-tufted, while neurons with ratios < 1 were classified as thin-tufted. Spine counts were obtained from 2^nd^-order branches of apical and basal dendrites located close to the surface for better visibility. Spines were counted on stretches of approximately 100 µm, whereby all types of spines were taken into account and pooled for analysis. Parameters of dendritic morphology were analyzed using Neuroexplorer software (MBF Bioscience, USA). Dendritic complexity was calculated as the sum of the terminal orders and the number of terminals, multiplied with the ratio of the total dendritic length and the number of primary dendrites (Pillai et al. 2012).

### Statistical tests

Unless stated otherwise, the non-parametrical Wilcoxon signed rank test and Mann-Whitney tests were used for paired and unpaired comparisons, respectively. Pair-wise noise and spontaneous correlations were assessed using Pearson’s correlation coefficient. Correlations between two variables were evaluated using the non-parametric Spearman’s rank correlation test, unless stated otherwise. All statistical tests were two-sided and were performed with either XLSTAT (Addinsoft, USA) or Matlab (Mathworks, USA). Statistical tests are specified in the figure legends and p-values are indicated using asterisks: * < 0.05; ** < 0.01; *** < 0.001.

